# Optogenetic decoupling of ODC inhibition and degradation reveals a requirement of polyamine oscillation for cell cycle progression

**DOI:** 10.64898/2026.07.16.739060

**Authors:** Zunqin Zhang, Bing Wu, Kan Wu, Zhiyan Chen, Sen Liu

## Abstract

The dysregulation of polyamine homeostasis is a fundamental hallmark of both cancer and aging. Ornithine decarboxylase (ODC), the rate-limiting enzyme in polyamine synthesis, is regulated by the antizyme OAZ1 through a dual mechanism: inhibition of enzymatic activity and induction of proteasomal degradation via a C-terminal degron. In Bachmann-Bupp syndrome (BABS), mutations in this degron lead to ODC accumulation and toxic polyamine levels. However, the functional coupling between ODC inhibition and degradation remains poorly understood, and the current therapeutic intervention with DFMO target enzymatic activity without addressing protein accumulation. Here, we developed an optogenetic yeast model of BABS to decouple ODC inhibition from its degradation. By fusing a light-switchable LOV2-based module to a truncated, degron-less yeast ODC, we created a system where ODC degradation is controlled by light, independent of OAZ1 binding. We demonstrate that the loss of the ODC degron mimics the BABS cellular phenotype, characterized by increased polyamines, elevated ROS, and growth arrest. Crucially, while constant light-induced degradation is insufficient to rescue growth, oscillating ODC degradation at a 40-minute period restores polyamine homeostasis, cell cycle progression, and cellular growth. Our findings demonstrate that the inhibitory and degradation-inducing roles of OAZ1 can be decoupled and highlight the necessity of periodic polyamine oscillation for the cell cycle. Our work would be of great value for understanding the regulation mechanism of the polyamine metabolic network and establishing new targeted therapeutic strategies.

**Significance statement:** Mutations in ODC degron lead to ODC accumulation and toxic polyamine levels in BABS. While DFMO has provided clinical relief for BABS patients, its utility is limited because it only inhibit ODC activity. We utilized an optogenetic approach in Saccharomyces cerevisiae to bridge this knowledge gap. We successfully decoupled the inhibitory and degradation-inducing functions of the ODC-OAZ1 axis and achieved three significant biological insights. First, we established yeast as a powerful and versatile model for investigating of the molecular mechanisms of BABS. Second, we demonstrated that the inhibitory and degradation-inducing roles of OAZ1 can be decoupled under specific conditions. Third, we revealed that periodic oscillation in polyamine levels is essential for maintaining proper cell cycle progressing.

## Introduction

Metabolic dysfunction is associated with various diseases. For instance, the three major polyamines in mammalian cells - putrescine, spermidine, and spermine - have indispensable biological functions, and the dysregulation of polyamine metabolism is associated with various abnormal cellular states ^1–3^. It has been widely observed that in cancer cells and tumor microenvironments, polyamine levels are much higher than in normal cells and organisms ^4–6^. Conversely, cellular polyamine content usually decreases in aging cells ^7–10^. Moreover, imbalances in the ratios of these polyamines also leads to diseases ^11,12^. In these abnormal states, rebalancing polyamine levels and ratios has been shown to be effective in recovering normal cellular functions ^9,11–14^. Recently, the dysregulation of the polyamine metabolic network has been recognized as a representative meta-hallmark of both aging and cancer ^15^.

The polyamine metabolic network consists of polyamine synthesis, metabolism, and membrane transport. ODC (ornithine decarboxylase 1) is the rate-limiting enzyme of the polyamine synthesis pathway; it catalyzes the decarboxylation of ornithine to generate putrescine, the precursor of spermidine and spermine. Through a feedback mechanism of +1 ribosomal frameshifting ^16^, polyamines stimulate the translation efficiency of OAZ1 (ODC antizyme 1) mRNA, a protein inhibitor of ODC. Upon binding to ODC, OAZ1 not only inhibits its enzymatic activity, but also accelerates its 26S proteasome dependent degradation by inducing the exposure of an unstructured sequence known as the “PEST degron” in ODC ^17^. Therefore, OAZ1-induced degradation exaggerates OAZ1’s inhibitory effect on ODC’s activity. After the degradation of the OAZ1-bound ODC, OAZ1 is recycled to replenish the OAZ1 pool.

As a key feedback mechanism regulating ODC activity, the inhibitory and the degradation-inducing roles of OAZ1 are tightly coupled, which may be indispensable for sustaining polyamine homeostasis. Indeed, mutations in the ODC degron sequence that disrupt its OAZ1-induced degradation cause various abnormalities, exemplified by Bachmann-Bupp syndrome (BABS) in human ^18–20^. In BABS, polyamine levels are abnormally elevated, and DFMO (α-difluoromethylornithine), an ODC inhibitor, has been used to treat this disease effectively ^19^. This demonstrates that the degron-dependent degradation of ODC is critical for normal cellular functions. However, although DFMO inhibits ODC’s enzymatic activity, it does not restore the degron-dependent degradation of ODC by the 26S proteasome. How the coupling of OAZ1-mediated inhibition and degradation of ODC contributes to the onset and treatment of BABS has not been systematically studied.

Polyamines are indispensable for virtually all cells, and many of their biological functions are conserved across organisms. For example, polyamine levels decrease during aging, and spermidine supplement has been shown to slow the aging process in humans and various model organisms including worms, yeast, flies, and mice ^21^. Due to its easy of genetic manipulation, low cost, and fast turnover, yeast is a valuable model organism for investigating human disease mechanisms, identifying drug targets, and discovering drug candidates. For instance, yeast has been a major tool for identifying spermidine as an anti-aging compound and elucidating its potential mechanism ^21,22^. Therefore, we reasoned that studying the cellular mechanism of BABS and the polyamine metabolic network in yeast could provide significant insights, as ODC-degradation-related diseases have not been investigated in this model yet.

In this work, we set out to determine how ODC degradation is coupled with its inhibition by OAZ1. To facilitate the experimental manipulation, we used budding yeast (*Saccharomyces cerevisiae*) as a model organism. To decouple the inhibitory function and the degradation-inducing function of the yeast OAZ1 (yOAZ1) on yODC (SPE1, referred to as yODC), and to make the decoupling process reversible, we adopted an optogenetic strategy. First, we validated that yeast exhibits cellular phenotypes similar to BABS when the ODC degron is missing. We then designed a yODC that retains its yOAZ1-inhibiting ability while gaining a light-switchable proteasome-dependent degradation capability. By incorporating this light-switchable yODC gene in the yeast chromosome, we demonstrated that light regulates both yODC degradation and fluctuations in polyamine concentration. We further showed that the yODC-inhibition function and the yODC-degradation-inducing function of yOAZ1 are decouplable when yODC degradation oscillates at a specific period. Finally, we demonstrated that the periodical yODC degradation facilitates cell cycle progression.

## Results

### 1. Degron removal decouples yODC’s yOAZ1-induced degradation from yOAZ1 inhibition

The polyamine metabolic network and the regulation mechanism of the yODC-yOAZ1 interaction in yeast are largely similar to those in human cells ^23^. A difference between yODC and human ODC (hODC) is that the degron sequence of yODC is located at the N-terminus, whereas it is located at the C-terminus in hODC. Previous studies showed that deleting the first 20 or 47 N-terminal residues slowed the yOAZ1-induced degradation of yODC ^24,25^. Therefore, we generated these two yODC truncations to determine how effectively they could decouple yODC’s yOAZ1-induced degradation from its other functions (enzymatic activity and yOAZ1 inhibition) (Figure 1a). We confirmed that the deletion of the N-terminal 20 (referred to as yODC_dN20_ for simplicity) or 47 (referred to as yODC_dN47_ for simplicity) residues did not compromise enzymatic activity (Figure 1b, Supplementary Fig. 1). The inhibition of their activity by yOAZ1 also remained comparable to the full-length wildtype yODC (Figure 1c, Supplementary Fig. 2). However, compared to full-length yODC (referred to as yODC for simiplicity), the mutants exhibited slower rates of spontaneous proteasomal degradation in the absence of yOAZ1, and yODC_dN47_ showed slower degradation than yODC_dN20_ (Figure 1d, Supplementary Fig. 3). In the presence of yOAZ1, while the degradation rates of yODC and yODC_dN20_ were similarly increased, yODC_dN47_ became more stable (Figure 1e, Supplementary Fig. 3), presumably due to a stabilization effect caused by protein-protein interactions. Note that yODC_dN20_ and yODC_dN47_ were expressed with a C-terminal 6xHis tag and an N-terminal FLAG tag (Supplementary Table 1), but our data indicate that these tags did not affect their function.

**Figure 1.**
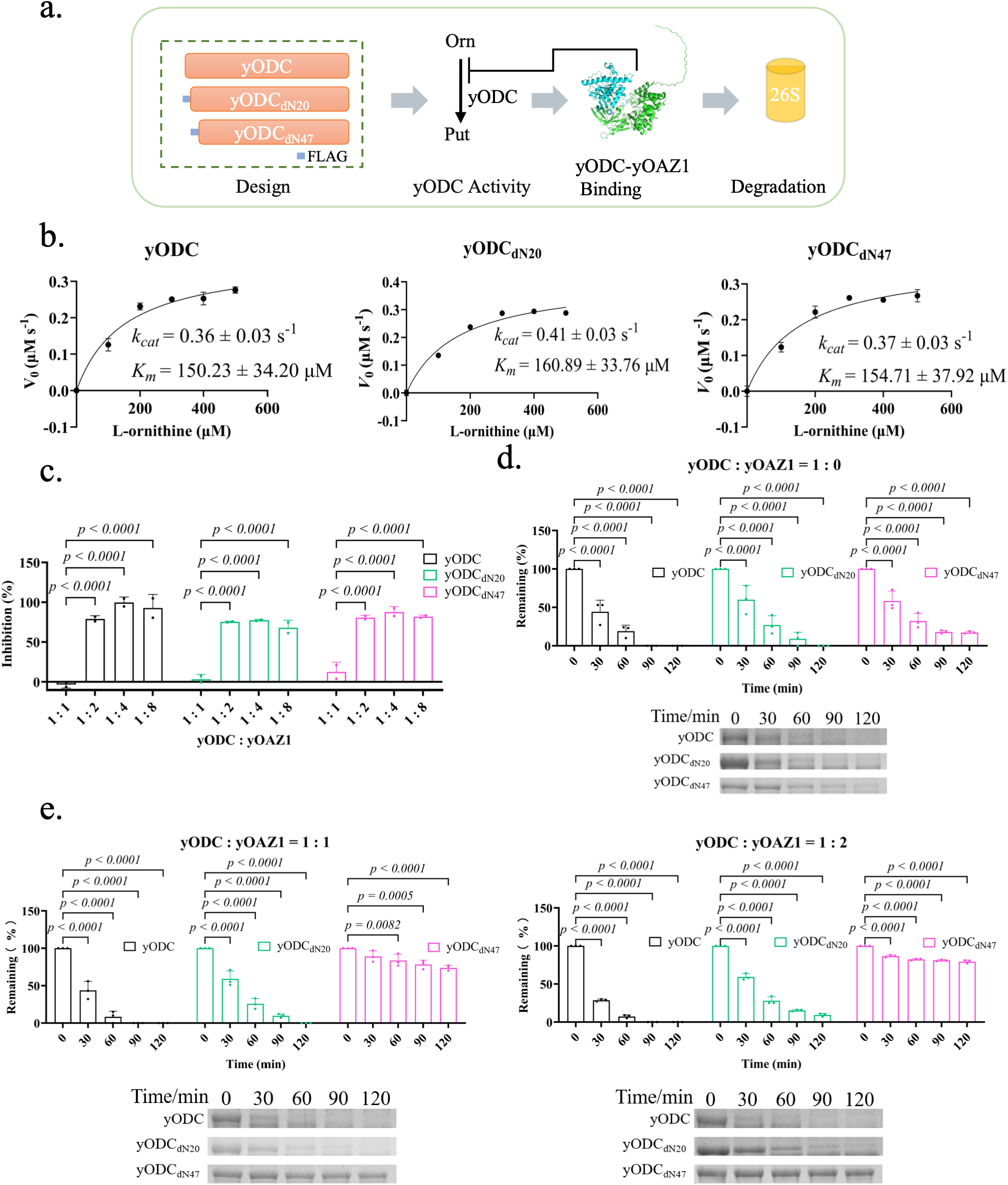
Degron removal decouples yODC’s yOAZ1-induced degradation from yOAZ1 inhibition. (a) A scheme of the experimental design for this section. (b) Enzymatic activity of different yODC constructs measured using a CO_2_ detection kit (Supplementary Fig. 1). (c) Inhibition of the enzymatic activity of different yODC constructs by yOAZ1 (Supplementary Fig. 2). (d, e) Proteasome-dependent degradation of the yODC proteins without (d) or with (e) yOAZ1. Representative protein bands are quantified using ImageJ. Full gels are provided in Supplementary Fig. 3.

Collectively, our data demonstrate that the removal of the N-terminal 20 or 47 degron residues of yODC decoupled its yOAZ1-induced degradation from its other functions. Since yODC_dN47_ was more stable than yODC_dN20_ against yOAZ1-induced degradation, we focused on this mutant for our subsequent investigations.

### 2. yODC_dN47_ leads to BABS-like cellular phenotypes in yeast

The clinical symptoms of BABS patients include developmental delay, hypotonia, non-congenital alopecia, and macrocephaly ^1^, accompanied by cellular phenotype changes such as ODC accumulation, increased ODC activity, and elevated polyamine levels ^19,26^. The ODC inhibitor DFMO is effective in treating BABS patients by reducing cellular ODC activity and polyamine levels ^19,26^. To investigate whether yeast exhibits BABS-like cellular phenotypes upon the removal of the yODC degron, we knocked out the wildtype yODC gene and knocked in the yODC_dN47_ gene in the *Saccharomyces cerevisiae* yeast strain W303 (Figure 2a). Yeast cells lacking the wild-type yODC gene (referred to as W303-ΔODC) exhibited minimal growth on SD (Synthetic Defined) medium (Figure 2b, Supplementary Table 2). While growth was recovered in the yODC_dN47_ strain (referred to as W303-FLAG-yODC_dN47_), the growth rate was significantly slower than the wildtype strain (referred to as W303-WT) (Figure 2b). Interestingly, while increased polyamine levels typically promote cell proliferation – as seen in tumor cells and fast-growing cells – this result aligns with a previous BABS study ^26^, which reported that the proliferation rate of primary dermal fibroblasts from an 11-month patient was much slower than that of normal neonatal primary dermal fibroblasts. Western blot analysis showed that yODC_dN47_ in W303-FLAG-yODC_dN47_ remained largely unchanged (Figure 2c, Supplementary Fig. 4). As expected, W303-FLAG-yODC_dN47_ had much higher levels of putrescine and spermidine than the wildtype strain (Figure 2d, Supplementary Fig. 5). This polyamine profile in W303-FLAG-yODC_dN47_ is consistent with changes observed in BABS patient cells ^26^, where putrescine and spermidine increased while spermine levels remained normal. When putrescine was added to the SD medium, the growth rate of W303-WT decreased (Figure 2e), confirming that high polyamine levels inhibit yeast growth. Because spermine levels were near our detection limit and appeared unresponsive to ODC alterations, we focused our subsequence polyamine analyses on putrescine and spermidine.

**Figure 2.**
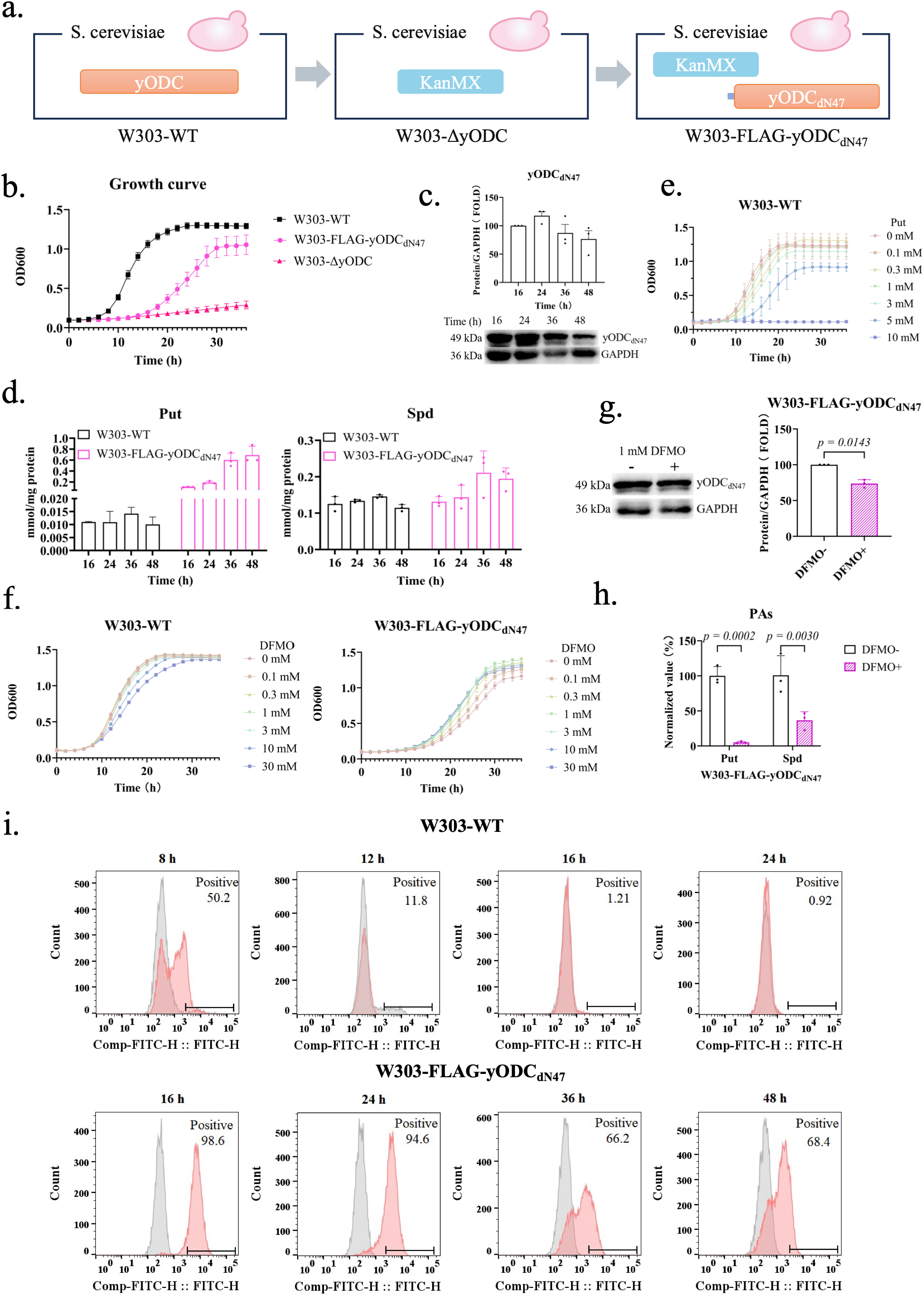
yODC_dN47_ leads to BABS-like cellular phenotypes in yeast. (a) A scheme of the W303 yeast strains used in this section. The KanMX cassette replaced the wild-type yODC gene, and the FLAG-yODC_dN47_ gene was integrated via *LEU2* homologous recombination. (b) Growth curves of different yeast strains in liquid SD medium. (c) yODC_dN47_ protein levels in W303-yODC_dN47_. The full western blot is shown in Supplementary Fig. 4. Bands were quantified using ImageJ. (d) Polyamine (putrescine and spermidine) contents in different yeast strains. (e) Growth curves of the wildtype W303 yeast strain in SD medium containing varying concentrations of putrescine. (f) Growth curves of different yeast strains under varying concentrations of DFMO in the liquid SD medium. (g) Representative western blot of yODC_dN47_ protein in W303-yODC_dN47_ upon DFMO treatment. The full blot is shown in Supplementary Fig. 6. Bands were quantified using ImageJ. (h) Polyamine contents in W303-yODC_dN47_ upon DFMO treatment. (i) Reactive oxygen species (ROS) levels in W303-WT and W303-yODC_dN47_. Grey peaks represent unstained negative controls, and red peaks indicate stained samples.

Since the accumulation of yODC and polyamines inhibits cell growth, we tested whether DFMO could reverse this growth arrest. Upon the addition of DFMO to the SD medium, the growth of W303-WT was slowed (Figure 2f); conversely, DFMO increased the growth rate of W303-FLAG-yODC_dN47_ (Figure 2f). Western blot analysis revealed that while DFMO treatment slightly decreased yODC_dN47_ protein levels (Figure 2g, Supplementary Fig. 6), it significantly reduced putrescine and spermidine concentrations (Figure 2h, Supplementary Fig. 7). Again, these results mirrored those of BABS patient cells ^26^, where DFMO treatment downregulated putrescine and spermidine (but not spermine). Additionally, W303-FLAG-yODC_dN47_ exhibited much higher reactive oxygen species (ROS) levels than the wild-type strain (Figure 2i), consistent with previous studies showing that elevated polyamine levels lead to increased ROS production ^27,28^.

Taken together, these data demonstrate that W303-FLAG-yODC_dN47_ exhibits phenotype changes similar to BABS patient cells ^26^, including slowed growth rates, accumulated ODC proteins, elevated polyamine levels, and effective response to DFMO. This similarity indicates that the regulation of the polyamine metabolic network is largely conserved between yeast and human cells, establishing yeast as a viable model organism for studying the cellular mechanism of BABS.

### 3. Light switches the proteasome-dependent degradation of yODC_dN47_-PSD3

When DFMO is used to treat BABS patients, it only inhibits ODC enzymatic activity and does not recover ODC’s OAZ1-induced proteasomal degradation. Furthermore, while the cellular levels of ODC and polyamines are different at different cell cycle phases ^29^, it is difficult to adjust cellular DFMO concentrations in a timely, reversible, or periodical manner. Optogenetics offers a fast-acting technique for the reversible regulation of protein function, allowing for precise temporal and periodic modulation. Therefore, we set out to design a yODC_dN47_ protein featuring a light-switchable module that regulates proteasomal degradation independent of yOAZ1 inhibition. We fused PSD3 ^30^, an optimized photo-switchable LOV2 construct containing the C-terminal 47 residues (degron) of hODC, to the C-terminus of yODC_dN47_ (Figure 3a, Supplementary Table 1). In darkness, the hODC degron is sequestered by the LOV2 core domain; upon blue light excitation, the degron is released and recognized by the proteasome, leading to the degradation of PSD3 and its fused target ^30^. First, we verified that in darkness, yODC_dN47_-PSD3 exhibited catalytic activity (Figure 3b, Supplementary Fig. 8) and yOAZ1 inhibition (Figure 3c, Supplementary Fig. 9) comparable to those of yODC_dN47_ and the wild-type yODC. Subsequent light recovery experiment confirmed that the PSD3 domain functioned normally within the fused construct (Figure 3d, Supplementary Fig. 10). Furthermore, upon light excitation, the presence of the PSD3 module did not affect the enzymatic activity (Figure 3e, Supplementary Fig. 11) or the OAZ1 inhibition (Figure 3f, Supplementary Fig. 12) of yODC_dN47_-PSD3. Proteasomal degradation assays showed that in darkness, the degradation of yODC_dN47_-PSD3 was similar to that of wild-type yODC and yODC_dN47_ (Figure 3g, Supplementary Fig. 13). As with yODC_dN47_ (Figure 1d), the degradation of yODC_dN47_-PSD3 was inhibited by MG132 (Figure 3g, Supplementary Fig. 13); however, yODC_dN47_-PSD3 exhibited a slightly higher degradation rate, likely due to the spontaneous release of the degron ^31,32^. Finally, we demonstrated that the degradation of yODC_dN47_-PSD3 was accelerated by light excitation and this acceleration was independent of yOAZ1 inhibition (Figure 3h, Supplementary Fig. 14).

**Figure 3.**
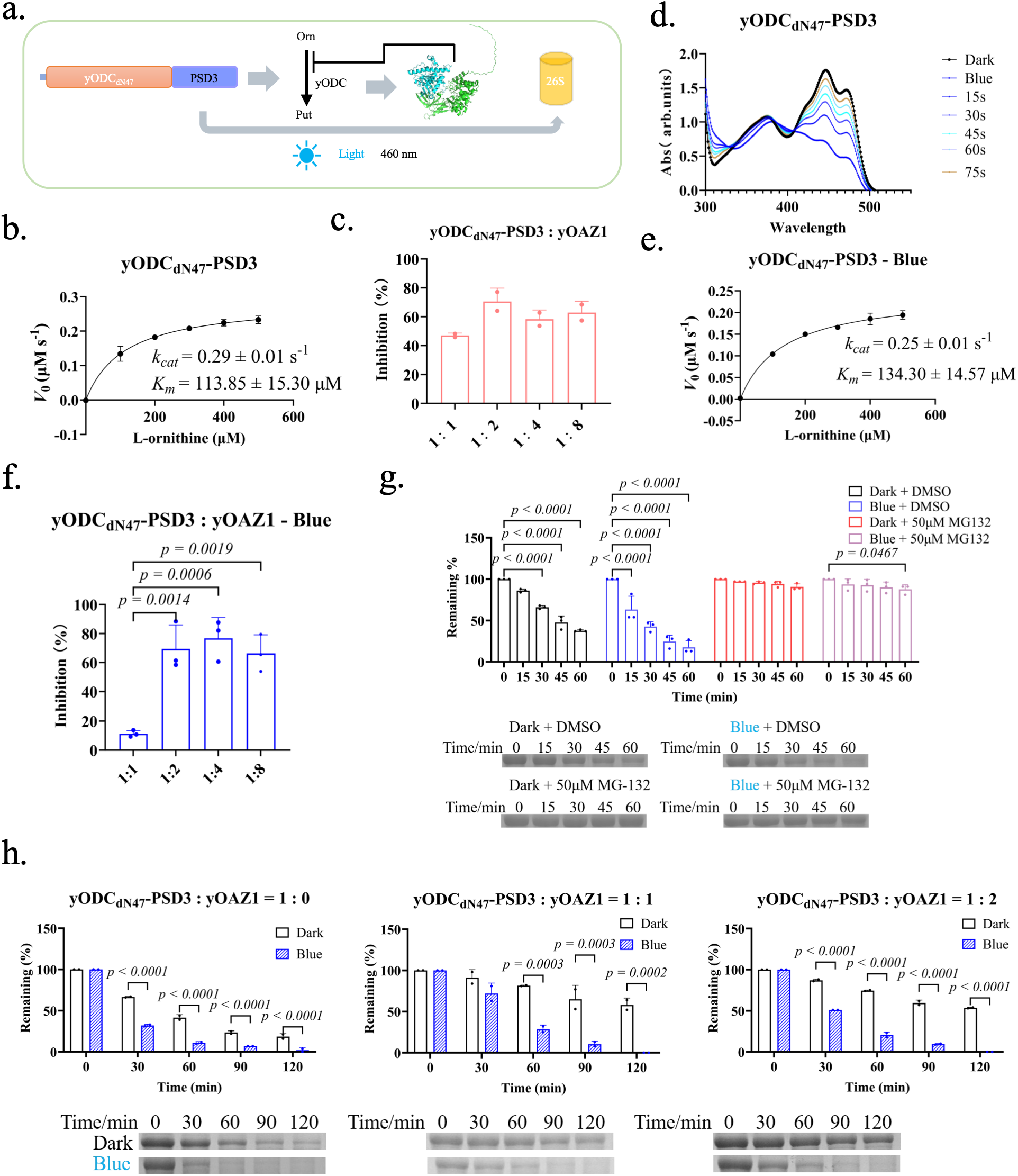
Light switches the proteasome-dependent degradation of yODC_dN47_-PSD3. (a) Schematic representation of the experimental design in this section. PSD3 is an optimized photo-switchable LOV2 construct containing the hODC C-terminal 47 residues (degron). The protein sequence of yODC_dN47_-PSD3 is shown in Supplementary Table 1. (b) Enzymatic activity of yODC_dN47_-PSD3 measured via CO_2_ detection (Supplementary Fig. 8). (c) Inhibition of yODC_dN47_-PSD3 enzymatic activity by yOAZ1 (Supplementary Fig. 9). (d) Change in light absorbance of yODC_dN47_-PSD3. (e) Enzymatic activity of yODC_dN47_-PSD3 upon light excitation measured via CO_2_ detection (Supplementary Fig. 11). (f) Inhibition of yODC_dN47_-PSD3 enzymatic activity by OAZ1 upon light excitation (Supplementary Fig. 12). (g) Inhibition of yODCdN47-PSD3 degradation by the proteasome inhibitor MG132. Representative SDS-PAGE bands and corresponding quantification from three biological replicates are shown; full gels are provided in Supplementary Fig. 13. (h) Proteasomal degradation of yODC_dN47_-PSD3 with or without yOAZ1. Representative SDS-PAGE bands and corresponding quantification from two biological replicates are shown; full gels are provided in Supplementary Fig. 14.

Taken together, these data confirm that fusing PSD3 to yODC_dN47_ created a yODC_dN47_ construct (yODC_dN47_-PSD3) with light-inducible proteasomal degradation that is decoupled from yOAZ1 inhibition.

### 4. yODC_dN47_-PSD3 induces BABS-like cellular phenotypes in yeast under dark conditions

We next investigated whether yODC_dN47_-PSD3 would induce BABS-like phenotypes in yeast under dark conditions by constructing a yeast mutant strain (W303-FLAG-yODC_dN47_-PSD3) (Figure 4a). As with W303-FLAG-yODC_dN47_, the W303-FLAG-yODC_dN47_-PSD3 strain showed a significantly slower growth rate than the wildtype strain, though the growth rates between the two mutant strains were comparable (Figure 4b). While yODC_dN47_-PSD3 protein levels remained relatively stable (Figure 4c, Supplementary Fig. 15), they were slightly lower than those of yODC_dN47_ in W303-FLAG-yODC_dN47_ (Figure 2c); this may be due to the spontaneous excitation of the PSD3 domain even in darkness ^31,32^,. Similar to the yODC_dN47_ strain (W303-FLAG-yODC_dN47_), W303-FLAG-yODC_dN47_-PSD3 accumulated much higher polyamine levels than the wildtype strain (Figure 4d, Supplementary Fig. 16). Furthermore, DFMO treatment did not inhibit but rather stimulated the growth of W303-FLAG-yODC_dN47_-PSD3 (Figure 4e). While DFMO did not affect yODC_dN47_-PSD3 protein levels (Figure 4f, Supplementary Fig. 17), it significantly reduced cellular polyamine concentrations (Figure 4g, Supplementary Fig. 18). Finally, ROS levels in W303-FLAG-yODC_dN47_-PSD3 (Figure 4h) were much higher than those of the wild-type strain (Figure 2i); however, these levels were lower than those observed in W303-FLAG-yODC_dN47_ (Figure 2i), likely reflecting the slightly lower absolute polyamine levels in W303-FLAG-yODC_dN47_-PSD3 (Figure 4d).

**Figure 4.**
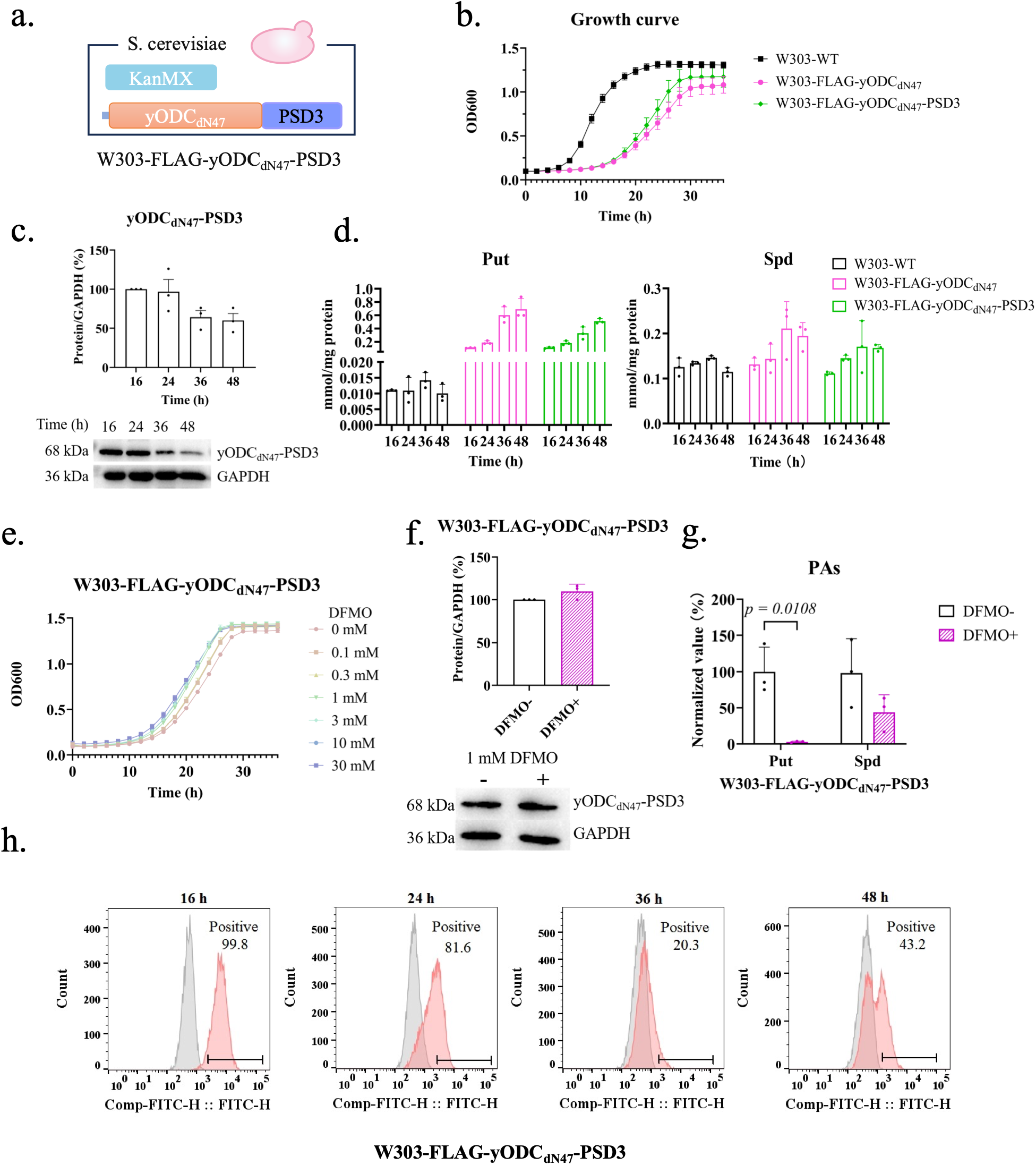
yODC_dN47_-PSD3 induces BABS-like cellular phenotypes in yeast under dark conditions. (a) A scheme of the W303 yeast strains used in this section. (b) Growth curves of different yeast strains in liquid SD medium. (c) yODC_dN47_-PSD3 protein levels in W303-FLAG-yODC_dN47_-PSD3. Representative western blot and corresponding quantification from three biological replicates are shown; the full blot is provided in Supplementary Fig. 15. (d) Polyamine contents in different yeast strains. (e) Growth curves of W303-FLAG-yODC_dN47_-PSD3 under varying concentrations of DFMO in liquid SD medium. (f) yODC_dN47_-PSD3 protein levels in W303-FLAG-yODC_dN47_-PSD3 upon DFMO treatment. Representative western blot and corresponding quantification from three biological replicates are shown; the full blot is provided in Supplementary Fig. 17. (g) Polyamine contents in W303-FLAG-yODC_dN47_-PSD3 upon DFMO treatment. Representative HPLC spectra are provided in Supplementary Fig. 18. (h) Reactive oxygen species (ROS) levels in W303-FLAG-yODC_dN47_-PSD3. Grey peaks represent unstained negative controls, and red peaks indicate stained samples.

Collectively, these data confirm that, much like the yODC_dN47_ strain (W303-FLAG-yODC_dN47_), the yODC_dN47_-PSD3 strain (W303-FLAG-yODC_dN47_-PSD3) exhibits BABS-like cellular phenotypes, although due to the intrinsic excitation of the LOV2 domain in the dark, the W303-FLAG-yODC_dN47_-PSD3 strain exhibited slightly lower levels of yODC_dN47_-PSD3 protein, polyamines, and ROS compared to the W303-FLAG-yODC_dN47_ strain.

### 5. Light regulates polyamine levels and yODC degradation in the W303-FLAG-yODC_dN47_-PSD3 strain

We next investigated whether the yODC_dN47_-PSD3 protein is responsive to light excitation in yeast. First, we observed that yODC_dN47_ levels in the W303-FLAG-yODC_dN47_ strain showed no significant difference under constant darkness, constant light, or during a light-darkness switch (Figure 5a, Supplementary Fig. 19). In contrast, yODC_dN47_-PSD3 levels in the W303-FLAG-yODC_dN47_-PSD3 strain significantly decreased under constant light and increased when the light was switched off (Figure 5b, Supplementary Fig. 20). Note that because these mutant strains exhibit slower growth rates than the wild-type strain, they were grown for 16 hours in darkness before undergoing light treatments. We then quantified polyamine levels to determine if cellular polyamines are responsive to light excitation. In W303-FLAG-yODC_dN47_, putrescine continued to accumulate under constant darkness, constant light, and during the light-darkness switch (Figure 5c, Supplementary Fig. 21). Conversely, in W303-FLAG-yODC_dN47_-PSD3, putrescine accumulated in constant darkness but decreased significantly under constant light; during the light-darkness switch, the putrescine levels were low but reversibly regulated (Figure 5d, Supplementary Fig. 22). Spermidine levels in both strains were lower than those of putrescine and showed no significant differences under varying light conditions (Figure 5b, Figure 5d, Supplementary Fig. 21, Supplementary Fig. 22). To confirm that the degradation of yODC_dN47_-PSD3 is light-induced and proteasome-dependent, we added the proteasome inhibitor MG132. While MG132 inhibited the degradation of yODC_dN47_-PSD3 under constant light (Figure 5e, Supplementary Fig. 23), the protein continued to accumulate whereas in constant darkness (Figure 5f, Supplementary Fig. 24). In contrast, yODC_dN47_ levels in W303-FLAG-yODC_dN47_ continued to accumulate under both constant darkness and constant light despite the presence of MG132 (Supplementary Fig. 25). Consistent with the changes in ODC and polyamines, ROS levels in the W303-FLAG-yODC_dN47_-PSD3 strain were lower under light than in darkness (Figure 5g, Figure 5h, Supplementary Fig. 26).

**Figure 5.**
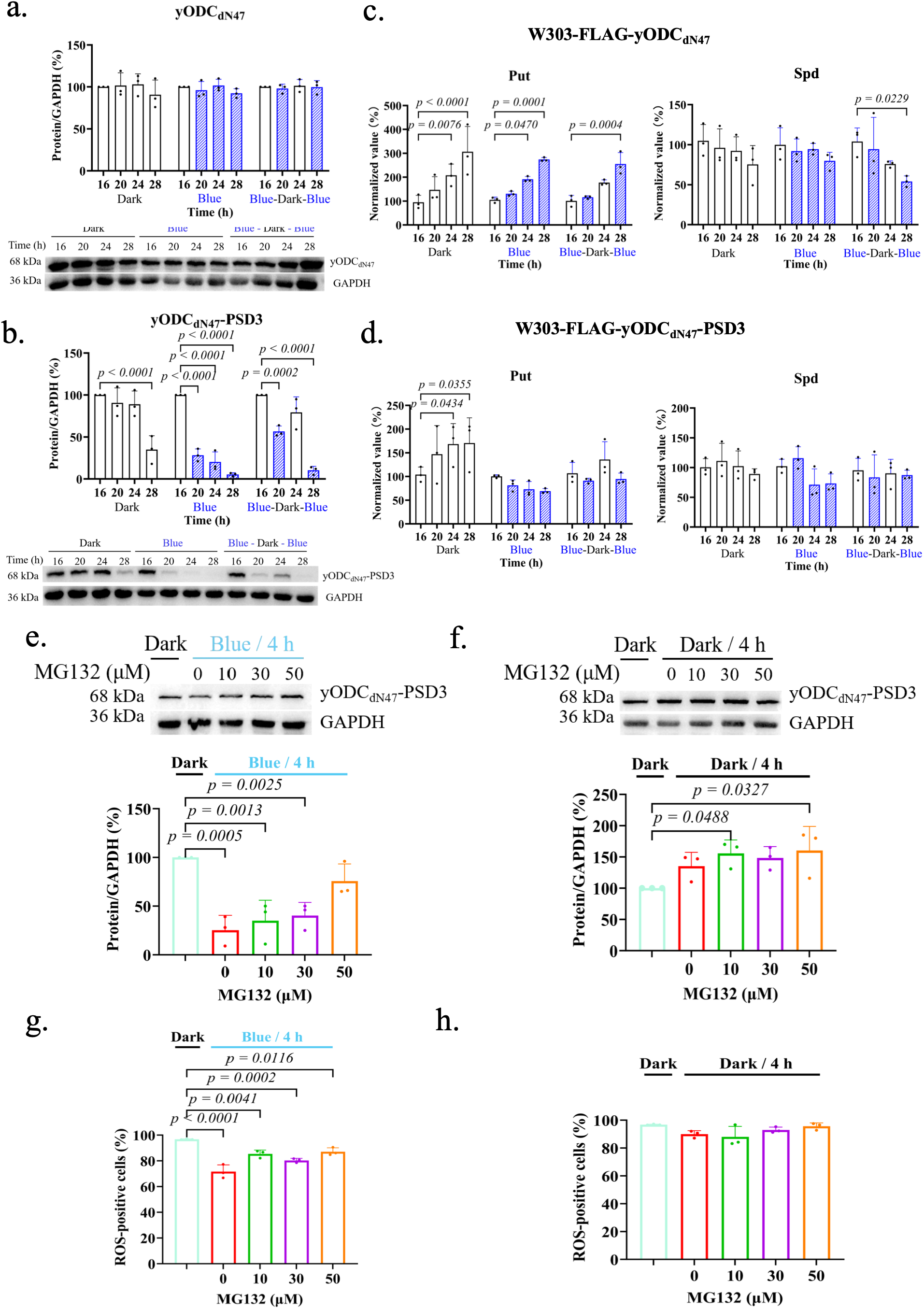
Light regulates polyamine levels and yODC degradation in the W303-FLAG-yODC_dN47_-PSD3 strain. (a) yODC_dN47_ protein levels in W303-FLAG-yODC_dN47_, which are not regulated by light. Representative western blot and corresponding quantification from three biological replicates are shown; the full blot is provided in Supplementary Fig. 19. (b) yODC_dN47_-PSD3 protein levels in W303-FLAG-yODC_dN47_-PSD3, which are regulated by light. Representative western blot and corresponding quantification from three biological replicates are shown; the full blot is provided in Supplementary Fig. 20. (c) Polyamine levels in W303-FLAG-yODC_dN47_, which are not regulated by light. (d) Polyamine levels in W303-FLAG-yODC_dN47_-PSD3, which are responsive to light. (e) Inhibition of light-induced proteosome-dependent degradation of yODC_dN47_-PSD3 by MG132 in W303-FLAG-yODC_dN47_-PSD3. Representative western blot and corresponding quantification from three biological replicates are shown; the full blot is provided in Supplementary Fig. 23. (f) Accumulation of yODC_dN47_-PSD3 in W303-FLAG-yODC_dN47_-PSD3 under constant darkness. Representative western blot and corresponding quantification from three biological replicates are shown; the full blot is provided in Supplementary Fig. 24. (g, h) ROS level changes in W303-FLAG-yODC_dN47_-PSD3 under light (g) and constant darkness (h).

Together, these data confirm that the degradation of yODC_dN47_-PSD3 can be reversibly regulated via light-dark switching in yeast and that this degradation is proteasome dependent. Furthermore, we demonstrated that this light-induced protein modulation results in corresponding, reversible changes in polyamines and ROS levels.

### 6. Periodic darkness/light oscillation rescues yeast growth arrest

Our findings demonstrated that polyamine accumulation, caused by the failure of yODC degradation, suppresses the growth of both W303-FLAG-yODC_dN47_ and W303-FLAG-yODC_dN47_-PSD3 in the dark; this growth arrest can be partially rescued by DFMO. For clinical treatment, besides the limitations mentioned earlier, persistent DFMO inhibition of ODC activity also induces undesirable adverse effects ^33^. Supporting this, consistent light exposure failed to rescue the growth of W303-FLAG-yODC_dN47_-PSD3 (Figure 6a, Supplementary Fig. 27). Given our ability to reversibly regulate yODC_dN47_-PSD3 degradation via optogenetics, we investigated whether darkness/light (DL) oscillation could rescue the growth arrest in the W303-FLAG-yODC_dN47_-PSD3 strain.

**Figure 6.**
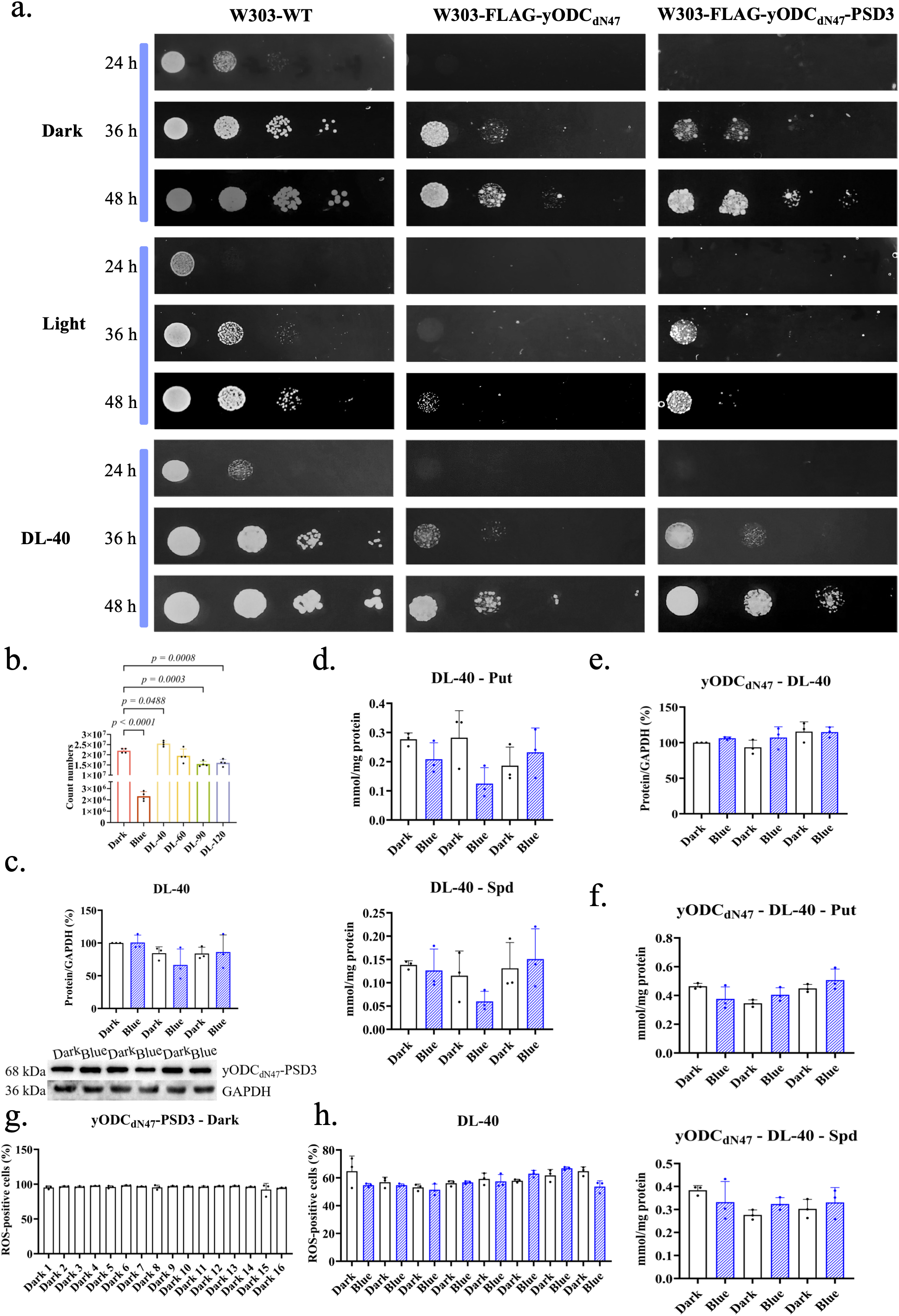
Periodic darkness/light oscillation rescues yeast growth arrest. (a) Growth of various yeast strains under different darkness/light (DL) conditions on SD plates. DL-40 indicates a 40-minute interval for blue light on/off switching. (b) Growth of W303-FLAG-yODC_dN47_-PSD3 under different DL conditions. (c) Changes in yODC_dN47_-PSD3 protein levels during DL-40 oscillation in W303-FLAG-yODC_dN47_-PSD3. Representative western blot and corresponding quantification from three biological replicates are shown; the full blot is provided in Supplementary Fig. 28. (d) Polyamine levels in W303-FLAG-yODC_dN47_-PSD3 during DL-40 oscillation. (e) yODC_dN47_ protein levels in W303-FLAG-yODC_dN47_ during DL-40 oscillation. (f) Polyamine levels in W303-FLAG-yODC_dN47_ during DL-40 oscillation. (g, h) ROS levels in W303-FLAG-yODC_dN47_-PSD3 under constant darkness (g) and DL-40 conditions (h).

Since ODC expression and polyamine synthesis fluctuate during cell cycle ^29^, and the typical doubling time of yeast is 90-120 min, we tested DL cycles with oscillation periods of 20 min (DL-20), 30 min (DL-30), 40 min (DL-40), and 60 min (DL-60). Under constant light, the growth of W303-WT was compromised, likely due to the toxicity of prolonged blue light exposure (Supplementary Fig. 27). Compared to constant darkness, the W303-FLAG-yODC_dN47_-PSD3 strain achieved improved growth rates under DL-40 (Figure 6a), whereas no further improvement was observed under DL-60 or extended oscillation periods (Figure 6a, Figure 6b, Supplementary Fig. 27). In contrast, the growth of W303-FLAG-yODC_dN47_ did not recover under these conditions (Figure 6a, Supplementary Fig. 27), confirming that the light-responsive PSD3 module is essential for this rescue. Noticeably, this 40-minute period is coincident with the DNA replication time of synchronized budding yeast ^34^. We further confirmed that both yODC_dN47_-PSD3 protein levels (Figure 6c, Supplementary Fig. 28) and polyamine levels (Figure 6d, Supplementary Fig. 29) in W303-FLAG-yODC_dN47_-PSD3 were reversibly modulated by the DL-40 oscillation. Conversely, in W303-FLAG-yODC_dN47_, the yODC_dN47_ protein level continued to accumulate (Figure 6e, Supplementary Fig. 30), and polyamine levels remained unresponsive and significantly elevated (Figure 6f, Supplementary Fig. 31). Finally, ROS level in W303-FLAG-yODC_dN47_-PSD3 were maintained at lower levels under DL-40 compared to constant darkness (Figure 6g, Figure 6h, Supplementary Fig. 32, Supplementary Fig. 33).

Taken together, these data indicate that the oscillation of yODC and polyamine levels at the 40-minute period facilitate cell cycle progression in W303-FLAG-yODC_dN47_-PSD3, likely by coupling with DNA replication and separation.

### 7. DL-40 and DFMO facilitate cell cycle progression but are non-additive

Since both DFMO treatment and DL-40 oscillation can regulate polyamine homeostasis in the W303-FLAG-yODC_dN47_-PSD3 strain, we investigated whether their combination could provide enhanced therapeutic benefits. Interestingly, although both DFMO and DL-40 oscillation independently improve the growth of W303-FLAG-yODC_dN47_-PSD3, their combination proved to be non-additive (Figure 7a, Figure 7b, Supplementary Fig. 34), confirming that they function in the same pathway. Under the combined treatment, the yODC_dN47_-PSD3 protein level in W303-FLAG-yODC_dN47_-PSD3 still responded to light modulation (Figure 7c, Supplementary Fig. 35); however, polyamine levels were less responsive than with DL-40 treatment alone, and absolute putrescine levels were significantly reduced (Figure 6c, Figure 7d, Supplementary Fig. 36). Despite this, the ROS levels remained low under the combined treatment (Figure 7e, Supplementary Fig. 37).

**Figure 7.**
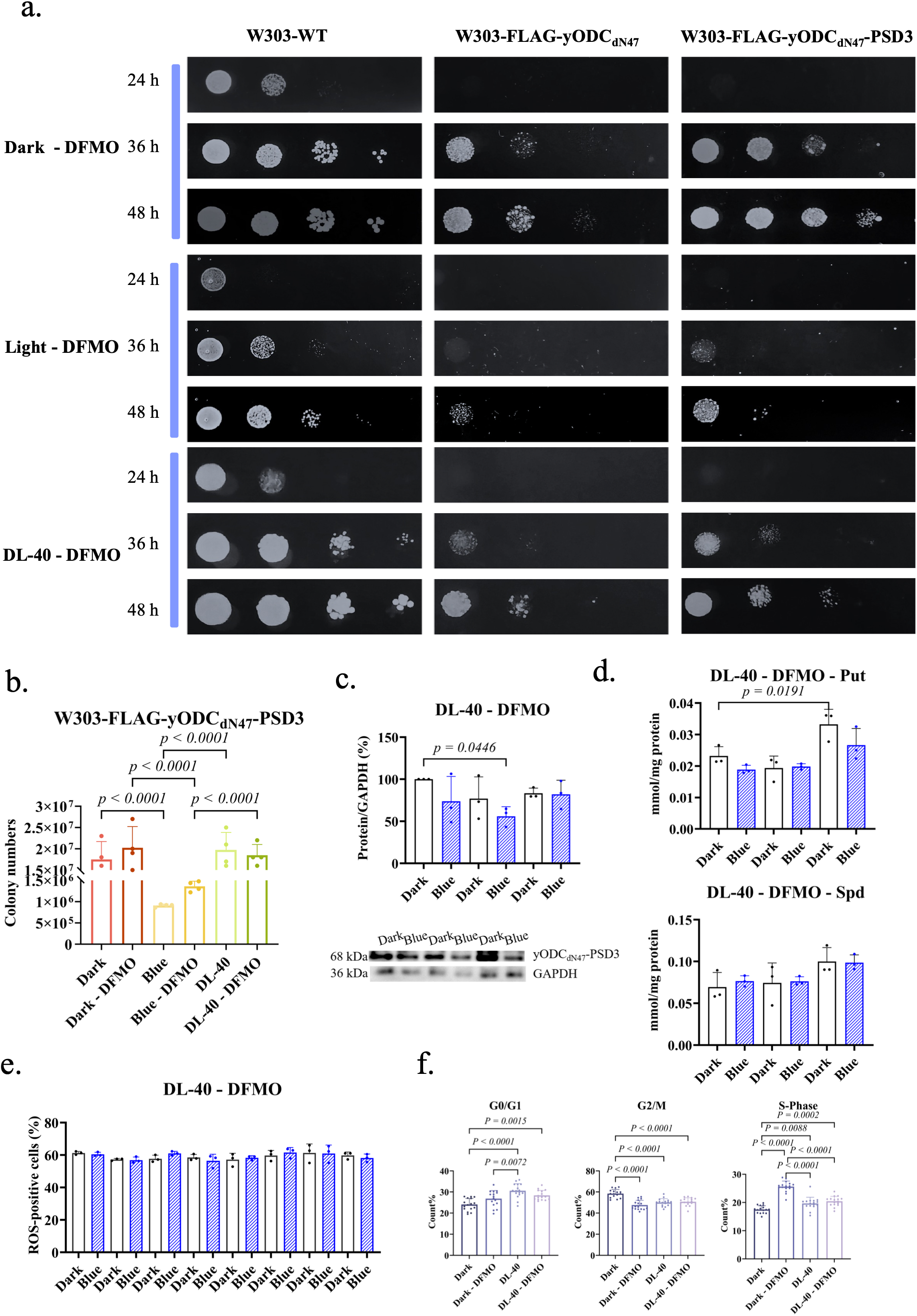
DL-40 and DFMO facilitate cell cycle progression but are non-additive. (a) Growth of various yeast strains on SD plates supplemented with 1mM DFMO under different light conditions. (b) Colony counts for W303-FLAG-yODC_dN47_-PSD3 under different conditions. (c) yODC_dN47_-PSD3 protein levels in W303-FLAG-yODC_dN47_-PSD3 during combined DL-40 and DFMO treatment. Representative western blot and corresponding quantification from three biological replicates are shown; the full blot is provided in Supplementary Fig. 35. (d) Polyamine levels in W303-FLAG-yODC_dN47_-PSD3 under combined DL-40 and DFMO treatment. (e) ROS positive cells in W303-FLAG-yODC_dN47_-PSD3 under combined DL-40 and DFMO treatment. (f) Cell cycle distribution of W303-FLAG-yODC_dN47_-PSD3 under different conditions. For each column, the data from 16 time points (Supplementary Fig. 38-S41) were averaged and are shown as mean ± s.e.m.

To investigate how the oscillations in ODC protein and cellular polyamine level affect cell cycle progression, we analyzed the cell cycle distribution of W303-FLAG-yODC_dN47_-PSD3 under single and combined DFMO and DL-40 treatments. The data revealed that upon treatments, significantly higher proportions of yeast cells proceeded to the G0/G1 phases compared to those in constant darkness (Figure 7f, Supplementary Fig. 38-S41). Meanwhile, the yeast cell exited the S and G2/M phases and entered the G0/G1 phase more often upon DL-40 treatment than DFMO treatment, indicating differences in underlying mechanism. The prolonged S phase under DFMO treatment agreed with previous findings on Chinese hamster ovary (CHO) cells ^35^ and human cells ^36,37^ under extended polyamine deprivation, revealing potential adverse effects of persistent DFMO treatment. The cell cycle distributions between DL-40 and the combined treatments were similar, indicating that DL-40 was the dominating factor. Accordingly, similar to DL-40 treatment, the combined treatment eliminated the prolonged S-phase caused by DFMO treatment alone.

Collectively, our data indicate that while both inhibiting yODC activity with DFMO and inducing oscillated yODC degradation are effective in reducing cellular polyamine levels and rescuing cell growth, their combination is redundant.

## Discussion and Conclusion

Polyaminopathies are a class of clinical diseases directly linked to the dysregulation of polyamine metabolism, with Bachmann-Bupp syndrome (BABS) being a prominent example due to its response to the FDA-approved ODC inhibitor DFMO ^1,18,38,39^. BABS originates from the truncation of the ODC C-terminal degron sequence, caused by premature stop codons resulting from gene mutations. The primary role of the ODC degron is to mediate ubiquitin-independent proteasomal degradation once ODC binds to and is inhibited by ODC antizymes (OAZs, primarily OAZ1). Consequently, truncation of this degron leads to the accumulation of both ODC protein and polyamines ^19,26,28^. While BABS patients showed beneficial responses to DFMO treatment, widespread clinical approval is currently hindered by limited case numbers and a lack of thorough mechanistic understanding. Because cellular and animal models are indispensable for studying of human disease mechanism – such as the K6/ODC mouse model, which mimics BABS phenotypes with a premature stop codon at position 427 of the mouse ODC ^28^ – the use of yeast as a powerful, cost-effective, and easily manipulatable single-cell model provides a vital complement to existing mammalian and animal models.

When the ODC C-terminal degron is partially or fully lost, a fundamental molecular consequence is the decoupling of OAZ1-mediated inhibition from OAZ1-induced degradation. Although these two functions are naturally coupled, whether they can be decoupled has remained uninvestigated. In this work, by leveraging optogenetics, we successfully demonstrated that these two functionalities of OAZ1 could be decoupled, at least partially, by oscillating ODC degradation at a 40-minute period synchronized with the cell cycle.

Currently, while DFMO treatment has shown promise for BABS patients, its application is limited. A significant drawback is that once administered, DFMO is difficult to remove or reverse, which is problematic for long-term use in human due to its well-identified side effects ^33^. Furthermore, while DFMO mimics the inhibitory effect of OAZ1 on ODC, it does not induce ODC degradation. Although our study did not observe additive benefits from combining DFMO treatment with light-induced ODC degradation, we demonstrated that inducing ODC degradation alone can achieve effects similar to or better than DFMO treatment. This finding suggests a novel therapeutic avenue for BABS: the use of ODC PROTACs (Proteolysis Targeting Chimeras) to actively induce ODC degradation.

Despite these findings, several limitations remain to be addressed. First, while our yeast model captures key characteristics of BABS and related mammalian cells, the full extent of its utility in representing human pathology requires further validation. Second, the effects of DFMO and DL oscillation were investigated under specific temporal and concentration parameters; whether optimized conditions could yield enhanced benefits remains to be determined. Finally, regarding the dual functions of OAZ1, our results raise a compelling new question: is OAZ1-induced proteasomal degradation of ODC a spontaneous process, or is it subject to additional regulatory mechanism within the cell?

In conclusion, this study has achieved three significant milestones. First, we established yeast as a powerful and versatile model for investigating of the molecular mechanisms of BABS. Second, we demonstrated that the inhibitory and degradation-inducing roles of OAZ1 can be decoupled under specific conditions. Third, we revealed that periodic oscillation in polyamine levels is essential for maintaining proper cell cycle progression. Collectively, these findings suggest that inducing ODC degradation may represent a highly effective therapeutic strategy for patients suffering from BABS.

## Methods

### 1. Materials

The *Saccharomyces cerevisiae* strains used in this study were derived from the W303 *MATa bar1* background, with the following genotype: *MATa ura3-1 his3-11 leu2-3 leu 2-112 trp1-1 can1-100 bar1Δ*, which was gifted by Professor Xiaojing Yang (Peking University). Plasmids pET-28a and pRS305 served as vectors for protein expression and yeast transformation/integration, respectively. Rabbit GAPDH polyclonal antibody (Cat. NO. 10494-1-AP) and rabbit DYKDDDDK tag polyclonal antibody (Cat. NO. 20543-1-AP) were from Proteintech (U.S.A). The HRP-conjugated goat anti-rabbit IgG (Cat. NO. D110058) was from Sangon Biotech (Shanghai) Co., Ltd., China. Putrescine (Cat. NO. V900377), spermidine (Cat. NO. S2626) and spermine (Cat. NO. S2876) were from Merck & Co. Inc., USA. MG132(Cat. NO. HY-13259), D,L-α-difluoromethylornithine (DFMO, Cat. NO. HY-B0744B) and isopropyl β-D-1-thiogalactopyranoside (IPTG, Cat. NO. A100487) were from MCE company (Shanghai), China. Carbon dioxide detection kit (Cat. NO. 100020260) was from Beijing Zhongsheng Baokuan Biotechnology Co., Ltd., China. 2’,7’-dichlorodihydrofluorescein diacetate (DCFH-DA, Cat. NO. D837204) was from Shanghai McLyn Biochemical Technology Co., Ltd., China. The BCA Protein quantification kit (Cat. NO. 20201ES76), homologous recombination cloning kit (Cat. NO. 10911ES20) and cell cycle detection kit (Cat. NO. 40301ES60) were from Yeasen Biotechnology (Shanghai) Co., Ltd., China. A 10 W, 460 nm blue-light lamp (Green Horizon Lighting Store) served as the light source. G418 was from Sangon Biotech (Shanghai) Co., Ltd., China. The 4.9 mW/cm² (Light intensity calculated as (Total electrical power × Electro-optical conversion efficiency) / (4π × Illumination distance²) was adopted as the optimal intensity preserving normal yeast growth.

### 2. Plasmid construction

The yeast ornithine decarboxylase (SPE1, referred to as yODC in this work) and the yeast ornithine decarboxylase antizyme 1 (yOAZ1) genes were amplified from yeast genomic DNA by PCR and individually cloned into pET-28a (digestion sites: BamHI and XhoI), with a 6×His tag fused to the C-terminus of yODC and the N-terminus of yOAZ1. The stop codon at the +1 frameshifting site of the yOAZ1 gene was eliminated by removing one nucleic acid (T) with overlap extension PCR for full-length yOAZ1 expression in *E. coli* ^40^. To assess protein expression in yeast cells with western blotting, the FLAG tag (DYKDDDDK) was added at the N-terminus of the ODC constructs (digestion site: Nco I). The PSD3 gene was synthesized by Tsingke Biotechnology (China). All plasmids were verified with DNA sequencing.

### 3. yODC deletion

The pADH1 promoter sequence was amplified by colony PCR. The amplified fragments were subsequently assembled with the KanMX resistance gene into pRS305 via homologous recombination. Homologous arms of 500 bp flanking the endogenous yODC locus were designed and fused to both ends of the KanMX gene via one-step cloning. The endogenous yODC gene was then replaced by the KanMX resistance cassette through homologous recombination (50°C, 30min). Yeast cells were cultured in YPD medium to mid-exponential phase, harvested by centrifugation at 2,500 rpm for 5 minutes, washed twice with sterile water, resuspended in 100 mM LiAc, and incubated at room temperature for 5 minutes. For the transformation, 240 μL of 50% PEG3350, 36 μL of 1 M LiAc, 5 μL of ssDNA, and 50 μL of linear DNA (pRS305) were sequentially added, followed by vortex mixing. Subsequently, 50 μL of competent cells were introduced into the mixture and incubated at 30 °C for 30 min. After adding 70 μL of DMSO, the cells were subjected to heat shock at 42 °C for 15 min. The cells were collected by centrifugation, resuspended in 1 mL of YPD medium, resuscitated at 30 °C for 1 hour, and then spread on plates for cultivation at 30 °C. Polyamine-deficient yeast strain W303-△yODC was selected on YPD plates supplemented with 700 μg/mL G418 and 1 mM Spd. Recombinant yeast strains W303-FLAG-yODC_dN47_ and W303-FLAG-yODC_dN47_-PSD3 were constructed via lithium acetate transformation as described previously ^41,42^.

### 4. Protein expression and purification

*E. coli* BL21(DE3) cells transformed with the indicated plasmids were cultured in LB medium supplemented with 50 μg/mL kanamycin at 37 °C to an OD₆₀₀ of 0.6-0.8, followed by induction with 0.5 mM IPTG at 16 °C for 16 h. Cells were harvested by centrifugation, resuspended in lysis buffer (50 mM Na_2_HPO₄, 150 mM NaCl, pH 7.5), disrupted by high-pressure homogenization, and centrifuged to remove debris. The supernatant was purified by Ni-NTA affinity chromatography to obtain His-tagged yODC and yOAZ1 proteins. Purified proteins were dialyzed and concentrated in storage buffer (50 mM Na_2_HPO₄, 150 mM NaCl, 1 mM DTT, 15% glycerol, pH 7.5), aliquoted, and stored at -80 °C.

### 5. Assay of yODC activity

The catalytic activity of yODC was determined using a CO_2_ detection kit, which monitors the decarboxylation of ornithine by measuring CO_2_ release to NADH oxidation as descripted previously ^6,43–45^. Briefly, yODC catalyzes the decarboxylation of ornithine, producing CO₂, which dissolves in the reaction buffer to form bicarbonate. Bicarbonate is then reacted with phosphoenolpyruvate (PEP) by phosphoenolpyruvate carboxylase (PEPC) to generate oxaloacetate, which is subsequently reduced to malate-by-malate dehydrogenase (MDH) with concurrent oxidation of NADH to NAD⁺. Since NADH absorbs light at 340 nm, the decrease in absorbance directly reflects yODC enzymatic activity. Assays were conducted in 96-well plates at 37°C. The substrate (ornithine) and cofactors (PEP and MgCl₂) were pre-incubated for 5 min with the enzyme mixture (1 μM yODC, PEPC, MDH, and NADH) before being mixed to initiate the reaction. NADH depletion was monitored kinetically at 340 nm for 10 min using a microplate reader. Initial velocities (V₀) were obtained by subtracting background absorbance, and the Michaelis constant (Kₘ) was determined by fitting the data to the Michaelis-Menten equation. For inhibition studies, purified yOAZ1 was incorporated into the reaction system, and the inhibitory effect was evaluated by measuring the relative decrease in NADH consumption rates over the first 5 min, normalized against blank (△A₀) and non-inhibited (△A₁) controls.

### 6. Assay of in vitro protein degradation

Wild-type yeast cells were cultured in YPD medium (Supplementary Table 2) until OD_600_ value reached approximately 15, and harvested by centrifugation at 3,000 rpm for 5 min at 4 °C. The cell pellet was resuspended in lysis buffer (25 mM Tris-HCl, pH 7.0, 10 mM MgCl_2_, 4 mM ATP,1 mM dithiothreitol (DTT), 10% glycerol (v/v)) and disrupted using a high-pressure homogenizer. The resulting lysate was centrifuged at 10,000 rpm for 15 min at 4 °C, and the supernatant (crude extract) was collected, aliquoted, and stored at -20 °C ^46^. Protein degradation assays were performed in a 250 μL reaction system containing 25 μL of the crude extract, 20 μg of the target protein, and 20 μL of degradation Buffer A (25 mM Tris-HCl, pH 7.0, 10 mM MgCl₂; 4 mM ATP and 1 mM DTT, added immediately before use), brought to the final volume with Buffer B (50 mM Na_2_HPO₄, 100 mM NaCl, 1 mM DTT, pH 7.0). After thorough mixing, the reaction was divided into 50 μL aliquots and incubated at 37 °C. At indicated time points (0, 30, 60, 90, and 120 min), reactions were terminated by adding 10 μL of 10× protein loading buffer followed by heating at 95 °C for 10 min. Samples were then resolved via 12% SDS-PAGE and visualized by Coomassie Brilliant Blue staining to evaluate degradation levels ^47^. For blue-light-induced degradation experiments, the reactions were performed under blue light irradiation (31.8 mW/cm²) at 37 °C. As of proteasome inhibition assays, crude extracts were pre-incubated with MG132 at 37°C for 15 min prior to addition to the reaction mixture under blue-light or dark conditions. Aliquots were pipetted out at 30-min intervals, and target protein degradation was monitored by SDS-PAGE.

### 7. Growth curve assay

For growth curve analysis, recombinant yeast was cultured overnight in SD medium to mid-exponential phase and diluted to OD₆₀₀ = 1.0. Aliquots (2 μL) were inoculated into 200 μL fresh SD medium in transparent flat-bottom 96-well plates, and growth was monitored continuously using a microplate reader (Bio-Tek Company, 5424R). To determine the effective concentration of DFMO on the strains, growth curves were measured in the presence of varying DFMO concentrations.

### 8. Western Blotting

Yeast cells were harvested and resuspended in sterile distilled water and lysed by vortexing with acid-washed glass beads (0.4-0.5 mm) for 1 min followed by 1 min on ice, repeated for 20 cycles ^48^. The lysate was clarified by centrifugation at 10,000 rpm for 15 min at 4°C, and the protein concentration was determined using the BCA assay. The cell lysate was subjected to SDS-PAGE electrophoresis, and the separated proteins were transferred onto a 0.22 μm PVDF membrane. The membrane was blocked with 5% non-fat milk for 2 h, incubated with primary antibody at 4 °C overnight, followed by incubation with secondary antibody at room temperature in dark for 2 h. Finally, protein bands were visualized by ECL chemiluminescence.

### 9. Determination of polyamine content

For polyamine derivatization, proteins in the lysed yeast samples above were precipitated with 0.2 M HClO₄. 500 μL of cell lysate was mixed with 20 μL of 20 mM DAH (internal standard), 500 μL of 2 M NaOH, and 10 μL of benzoyl chloride (derivatization reagent). The mixture was vortexed thoroughly and incubated at 40 °C for 20 min. After incubation, 2 mL of saturated NaCl solution was added and vortexed to stop the reaction. Then, 2 mL of dichloromethane was added for extraction; the lower organic phase was transferred to a new EP tube, and the extraction procedure was repeated three times. The organic (lower) phases were pooled, evaporated to dryness, and reconstituted in 200 μL of methanol. The solution was filtered through a 0.22 μm organic membrane filter, and 20 μL was injected for High-Performance Liquid Chromatography (HPLC, Thermo Fisher Scientific, DYY-6C) analysis. Separation was performed on a C18 column at 30°C with a flow rate of 0.5 mL/min using a gradient of 0.1% formic acid in water (mobile phase A) and 0.1% formic acid in acetonitrile (mobile phase B): 40% B from 0 to 4 min, increasing to 60% B from 4 to 35 min, and the absorbance was monitored at 254 nm ^49^.

### 10. Analysis of PSD3 function

2 mL of 50 μM FLAG-yODC_dN47_-PSD3 was transferred to a cuvette and irradiated with blue light for 5 min in dark or white-light background conditions, with images recorded at 10 s intervals. Full-wavelength scanning was then performed using a microvolume spectrophotometer: an initial dark-state spectrum was acquired, the sample was re-irradiated with blue light for 5 min, and spectra were subsequently recorded at 10 s intervals until the dark-state baseline was recovered.

### 11. Evaluation of cellular protein and polyamine changes

Recombinant yeast strains were cultured overnight in SD medium to mid-exponential phase, diluted to OD₆₀₀ = 1.0, and reinoculated into fresh SD medium. Cultures were incubated at 30 °C, 200 rpm, and cells were harvested at indicated time points for protein and polyamine quantification. To assess the impact of DFMO, cultures supplemented with 1 mM DFMO were maintained under dark conditions, and protein and polyamine levels were determined as described above.

### 12. Assessment of light-dependent degradation and reversibility of yODC in cell

Recombinant yeast cells entering the logarithmic phase were dark-adapted in SD medium for 16 h. Cultures were subsequently maintained in constant darkness, constant blue light, or blue light/darkness oscillation. Samples were taken every 4 h to assess temporal dynamics of protein and polyamine concentrations.

### 13. Evaluation of light-induced proteosome-dependent degradation of yODC in cell

Yeast cultures were grown overnight in SD medium to mid-exponential phase, diluted to OD600 = 1.0, and transferred to fresh medium. After 16 h of dark adaptation, samples (20 mL) were treated with DMSO or MG132 at indicated concentrations and incubated under blue light or darkness for 4 h. Cells were harvested, washed, and lysed by glass bead disruption. Protein concentrations were determined, and protein expression was analyzed by Western Blotting.

### 14. Effect of light oscillation on yeast strains

Yeast cells were grown overnight in SD medium to mid-exponential phase and diluted to OD₆₀₀ = 1.0. Aliquots (5 μL) of serially diluted cultures were spotted onto SD agar plates ^50^ and incubated under constant darkness, constant blue light, or dark/blue light oscillations (20-60 min). For protein and polyamine analyses, the diluted culture was inoculated into 200 mL of fresh SD medium and grown for 16 h under dark/blue light oscillation (40 min). Samples were collected starting from the onset of the first dark cycle for protein Western Blotting analysis and polyamine quantification. To investigate the combined effects of DFMO and dark/blue light oscillation on polyamine homeostasis, 1 mM DFMO was added to the aforementioned culture conditions.

### 15. Cellular ROS evaluation

Yest cells were harvested by centrifugation at 3,000 rpm for 5 min, washed twice with an equal volume of ultrapure water, and diluted to an OD₆₀₀ of 0.2. 1 mL of the diluted cell suspension was filtered through a 400-mesh sieve, followed by the addition of 10 μL 10 mM DCFH-DA. The mixture was gently inverted and incubated for 1 h in the dark. The fluorescence intensity was subsequently detected using a flow cytometer (Attue NxT) in the FITC channel (excitation, 488 nm; emission, 530 nm) ^51^.

### 16. Flow cytometric analysis of cell cycle

Yeast cells were harvested and adjusted to OD₆₀₀ = 0.1. Cells were pelleted by centrifugation at 3,000 rpm, 4 °C for 5 min, and the supernatant was discarded. The pellet was gently resuspended in 1 mL of ice-cold 70% ethanol and fixed at 4 °C for 2 h or overnight. After fixation, cells were collected by centrifugation at 3,000 rpm, 4 °C for 5 min, and washed twice with 1 mL of 50 mmol/L sodium citrate solution (3,000 rpm, 4 °C, 5 min each). For DNA staining, a working solution was prepared by adding 10 μL of propidium iodide (PI) and 10 μL of RNase A to 0.5 mL of staining buffer supplemented with RNase and PI from the cell cycle detection kit. Cells were resuspended in 0.5 mL of the working solution, incubated at 37 °C in dark for 30 min, and filtered through a 400-mesh sieve prior to analysis. Flow cytometry (Attue NxT) was performed in the PI channel (excitation, 488 nm; emission, 574 nm), with simultaneous detection of fluorescence and light scattering ^52^.

### 17. Statistical analyses

Statistical analyses were performed with GraphPad Prism (10.6.2), and the significance of the results was evaluated using the Student-Newman-Kuels test. The data were presented as mean ± standard deviation (s.d.) unless indicated otherwise. All assays were performed with at least two technical replicates and three biological replicates.

## Acknowledgements

We would like to thank the support from the other members of our lab. We thank Professor Xiaojing Yang for kindly providing us with the wildtype W303 yeast strain and suggestions.

## Author contributions

**Sen Liu**: Writing - review & editing, Writing - original draft, Supervision, Resources, Project administration, Funding acquisition, Formal analysis, Conceptualization. **Zunqing Zhang**: Writing - original draft, Visualization, Validation, Methodology, Investigation, Formal analysis. **Bing Wu**: Writing - original draft, Visualization, Validation, Methodology, Investigation, Formal analysis. **Kang Wu**: Validation, Methodology, Investigation. **Zhiyan Chen**: Visualization, Validation, Methodology, Investigation. All authors reviewed and approved the submitted manuscript.

## Data availability

Data and materials are available from the corresponding author upon reasonable request.

