## Supplemental Data for "Optogenetic decoupling of ODC inhibition and degradation reveals a requirement of polyamine oscillation for cell cycle progression"

**Supplementary Table 1**. The expressed sequences of the proteins mentioned in this paper.

| Protein | Sequence | Note |
|---|---|---|
| yODC | MSSTQVGNALSSSTTTLVDLSNSTVTQKKQYYKDGETLHNLLLE<br>LKNNQDLELLPHEQAHPKIFQALKARIGRINNETCDPGEENSFFIC<br>DLGEVKRLFNNWVKELPRIKPFYAVKCNPDTKVLSLLAELGVNF<br>DCASKVEIDRVLSMNISPDRIVYANPCKVASFIRYAASKNVMKSTF<br>DNVEELHKIKKFHPESQLLLRIATDDSTAQCRLSTKYGCEMENVD<br>VLLKAIKELGLNLAGVSFHVGSGASDFTSLYKAVRDARTVFDKA<br>ANEYGLPPLKILDVGGGFQFESFKESTAVLRLALEEFFPVGCGVDI<br>IAEPGRYFVATAFTLASHVIAKRKLSENEAMIYTNDGVYGNMNCI<br>LFDHQEPHPRTLYHNLEFHYDDFESTTAVLDSINKTRSEYPYKVSI<br>WGPTCDGLDCIAKEYYMKHDVIVGDWFYFPALGAYTSSAATQFN<br>GFEQTADIVYIDSELDGSSHHHHHH | Full-length wildtype yODC, containing a C-terminal 6xHis tag |
| yOAZ1 | MGSSHHHHHHSSGLVPRGSHMASMTGGQQMGRGSMYEVIQKR<br>KTKIINVLQSPELMRLIEDPSNLGISLHFPVSSLLKSNKCTPMPKLS<br>TYSLASGGFKDWCADIPLDVPPEIDIIDFYWDVILCMESQFILDYN<br>VPSKNKGNNQKSVAKLLKNKLVNDMKTTLKRLIYNENTKQYKN<br>NNSHDGYNWRKLGSQYFILYLPLFTQELIWCKLNENYFHVVLPS<br>LLNSRNVHDNHSTYINKDWLLALLELTSNLNQNFKFEYMKLRLY<br>ILRDDLINNGLDLLKNLNWVGGKLIKNEDREVLLNSTDLATDSIS<br>HLLGDENFVILEFEC | Full-length wildtype yOAZ1, containing an N-terminal 6xHis tag |
| yODC$_{dN20}$ | MDYKDDDDKLSNSTVTQKKQYYKDGETLHNLLLELKNNQDLEL<br>LPHEQAHPKIFQALKARIGRINNETCDPGEENSFFICDLGEVKRLF<br>NNWVKELPRIKPFYAVKCNPDTKVLSLLAELGVNFDCASKVEIDR<br>VLSMNISPDRIVYANPCKVASFIRYAASKNVMKSTFDNVEELHKIK<br>KFHPESQLLLRIATDDSTAQCRLSTKYGCEMENVDVLLKAIKELG<br>LNLAGVSFHVGSGASDFTSLYKAVRDARTVFDKAANEYGLPPLKI<br>LDVGGGFQFESFKESTAVLRLALEEFFPVGCGVDIIAEPGRYFVAT<br>AFTLASHVIAKRKLSENEAMIYTNDGVYGNMNCILFDHQEPHPR<br>TLYHNLEFHYDDFESTTAVLDSINKTRSEYPYKVSIWGPTCDGLD<br>CIAKEYYMKHDVIVGDWFYFPALGAYTSSAATQFNGFEQTADIV<br>YIDSELDGSSHHHHHH | yODC without the N-terminal 20 residues, containing an N-terminal FLAG tag and a C-terminal 6xHis tag |
| yODC$_{dN47}$ | MDYKDDDDKNQDLELLPHEQAHPKIFQALKARIGRINNETCDPG<br>EENSFFICDLGEVKRLFNNWVKELPRIKPFYAVKCNPDTKVLSLL<br>AELGVNFDCASKVEIDRVLSMNISPDRIVYANPCKVASFIRYAASK<br>NVMKSTFDNVEELHKIKKFHPESQLLLRIATDDSTAQCRLSTKYG<br>CEMENVDVLLKAIKELGLNLAGVSFHVGSGASDFTSLYKAVRDA<br>RTVFDKAANEYGLPPLKILDVGGGFQFESFKESTAVLRLALEEFFP<br>VGCGVDIIAEPGRYFVATAFTLASHVIAKRKLSENEAMIYTNDGV<br>YGNMNCILFDHQEPHPRTLYHNLEFHYDDFESTTAVLDSINKTRS<br>EYPYKVSIWGPTCDGLDCIAKEYYMKHDVIVGDWFYFPALGAYT<br>SSAATQFNGFEQTADIVYIDSELDGSSHHHHHH | yODC without the N-terminal 47 residues, containing an N-terminal FLAG tag and a C-terminal 6xHis tag |
| yODC$_{dN47}$-PSD3 | MGSSHHHHHHSSGHMGSLEVLFQGPDYKDDDDKNQDLELLPHE<br>QAHPKIFQALKARIGRINNETCDPGEENSFFICDLGEVKRLFNNW<br>VKELPRIKPFYAVKCNPDTKVLSLLAELGVNFDCASKVEIDRVLS<br>MNISPDRIVYANPCKVASFIRYAASKNVMKSTFDNVEELHKIKKF<br>HPESQLLLRIATDDSTAQCRLSTKYGCEMENVDVLLKAIKELGLN<br>LAGVSFHVGSGASDFTSLYKAVRDARTVFDKAANEYGLPPLKILD<br>VGGGFQFESFKESTAVLRLALEEFFPVGCGVDIIAEPGRYFVATAFT<br>LASHVIAKRKLSENEAMIYTNDGVYGNMNCILFDHQEPHPRTLY<br>HNLEFHYDDFESTTAVLDSINKTRSEYPYKVSIWGPTCDGLDCIA<br>KEYYMKHDVIVGDWFYFPALGAYTSSAATQFNGFEQTADIVYIDS<br>ELDGSRTSGGSGGSLATTLERIEKNFVITDPRLPDNPIIFASDSFLQL<br>TEYSREEILGRNCRFLQGPETDRATVRKIRDAIDNQTEVTVQLINY<br>TKSGKKFWNVFHLQPMRDYKGDVQYFIGVQLDGTERLHGAAER<br>EAVMLIKKTAFQIAEAKELPMSCAQESITSLYKKAGSENLYFQ | yODC without the N-terminal 47 residues, containing an N-terminal 6xHis/FLAG tag, containing a C-terminal PSD3 sequence |

**Supplementary Table 2**. The culture media used in this paper.

| Medium | Component | g/L (YPD) or mg/L (SD) |
|---|---|---|
| YPD | Yeast extract | 10 |
| | Peptone | 20 |
| | Glucose | 20 |
| SD (Synthetic Defined) | Yeast nitrogen base | 6700 |
| | Glucose | 20000 |
| | Adenine sulfate | 40 |
| | Arginine | 20 |
| | L-Aspartic acid | 100 |
| | L-Glutamic acid | 100 |
| | L-Histidine | 20 |
| | L-Lysine | 30 |
| | L-Methionine | 20 |
| | L-Phenylalanine | 50 |
| | L-Serine | 375 |
| | L-Threonine | 200 |
| | L-Tryptophan | 40 |
| | L-Tyrosine | 30 |
| | L-Valine | 150 |
| | L-Leucine | 100 |
| | Uracil | 20 |

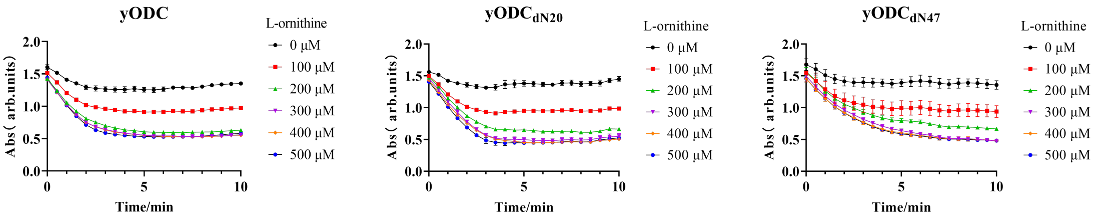


**Supplementary Figure 1**. Enzymatic activity data of different yODC constructs from the $CO_2$ detection kit.

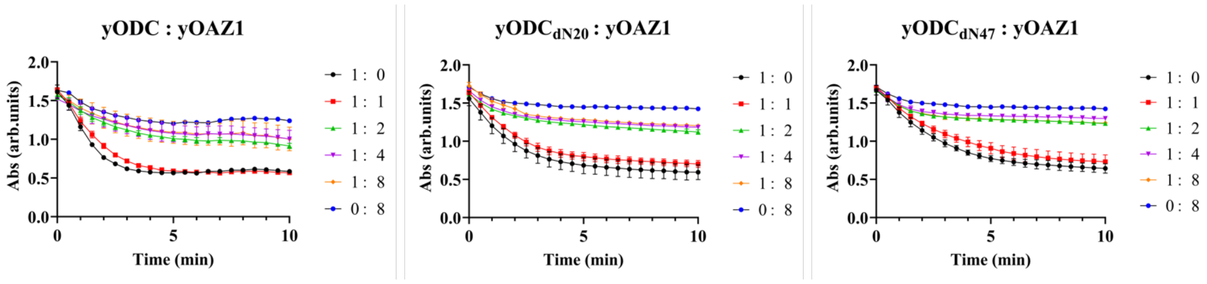


**Supplementary Figure 2**. Inhibition of yODC activity by yOAZ1. The molar ratios of yODC and yOAZ1 are shown in the legends.

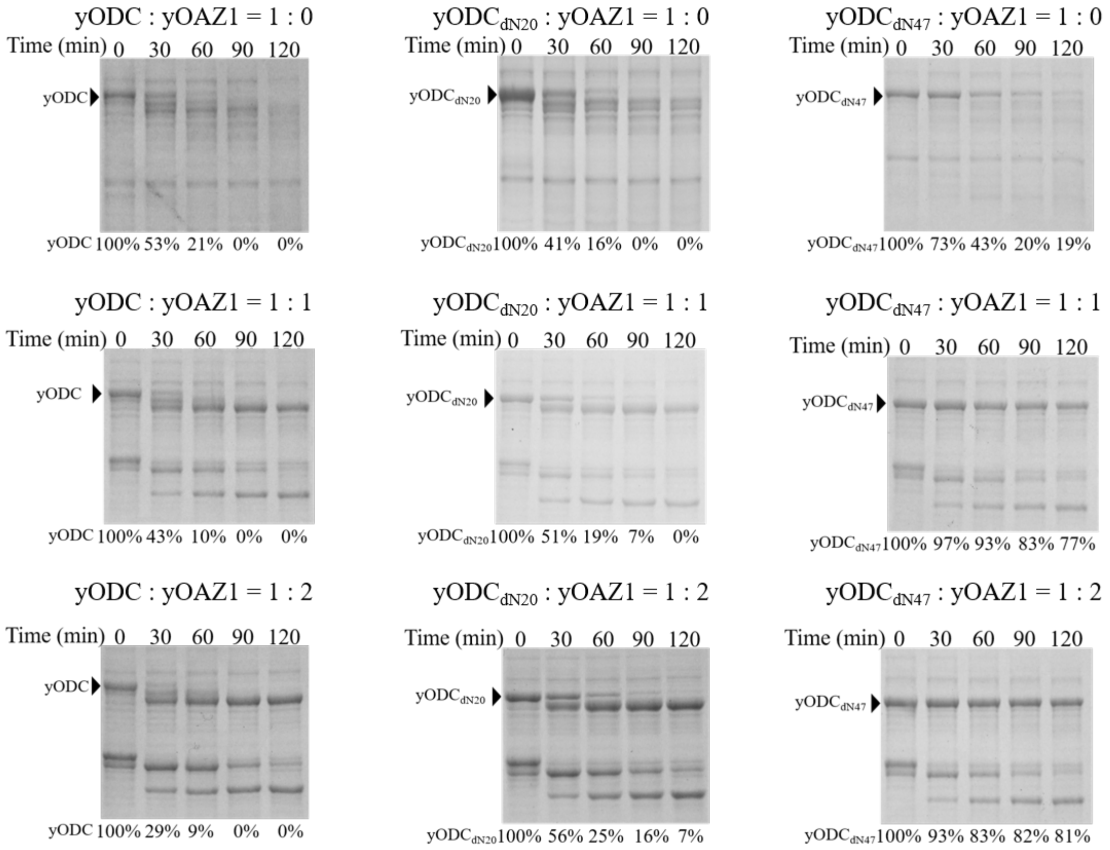


**Supplementary Figure 3**. The proteasome-dependent degradation of yODC constructs under different yODC-yOAZ1 ratios.

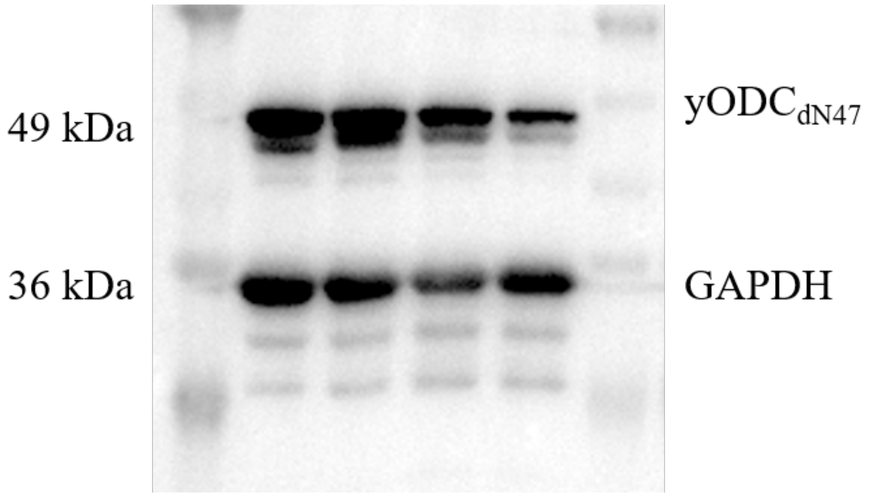


**Supplementary Figure 4**. The full western blotting gel of Figure 2c.

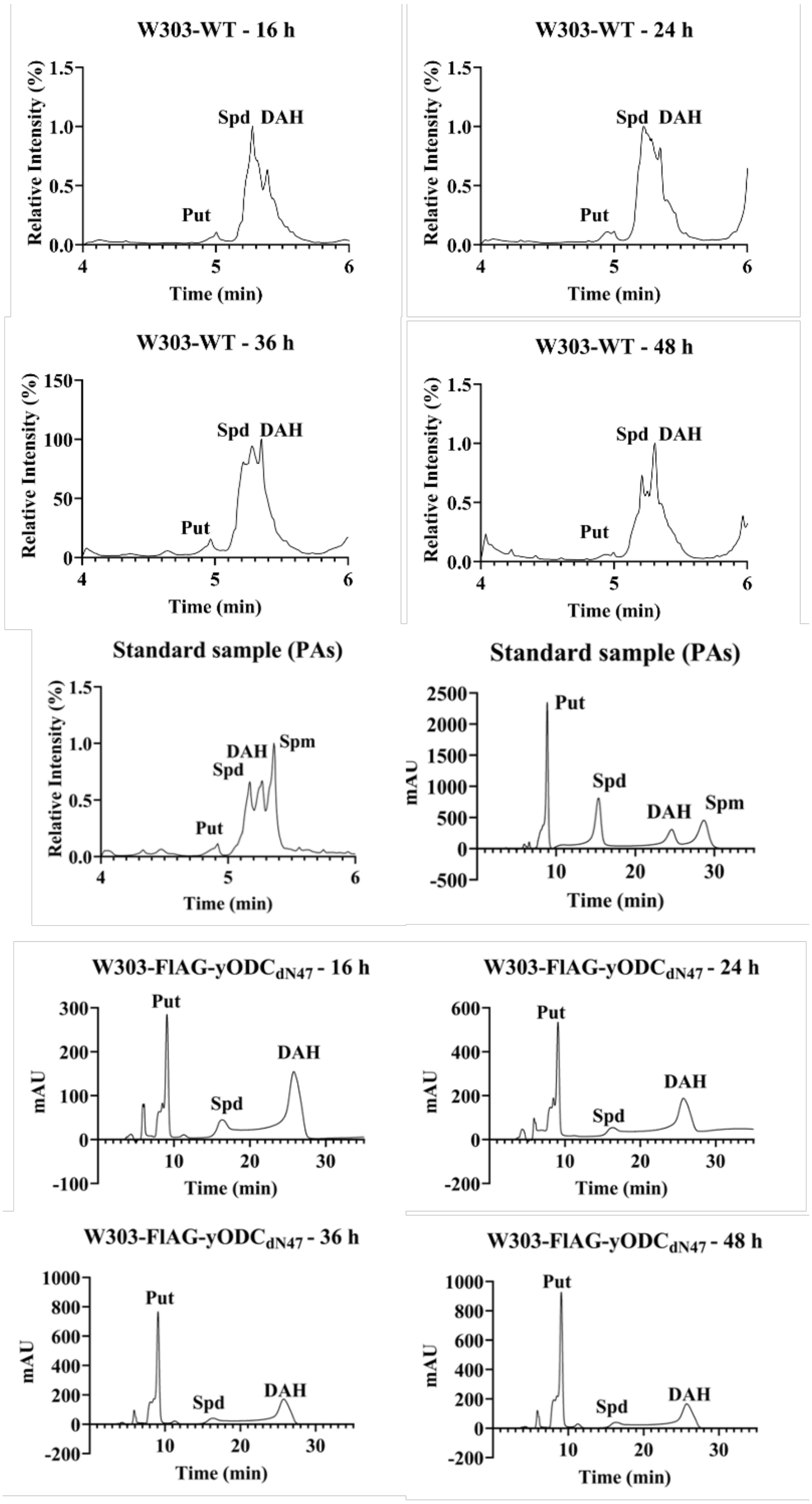


**Supplementary Figure 5**. Representative HPLC spectra for polyamine standards and samples of Figure 2d.

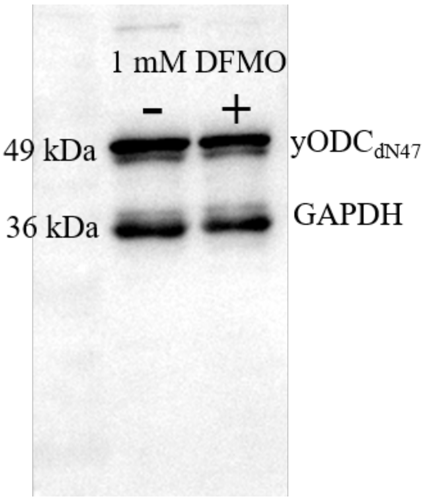


**Supplementary Figure 6**. The full western blotting gel of Figure 2g.

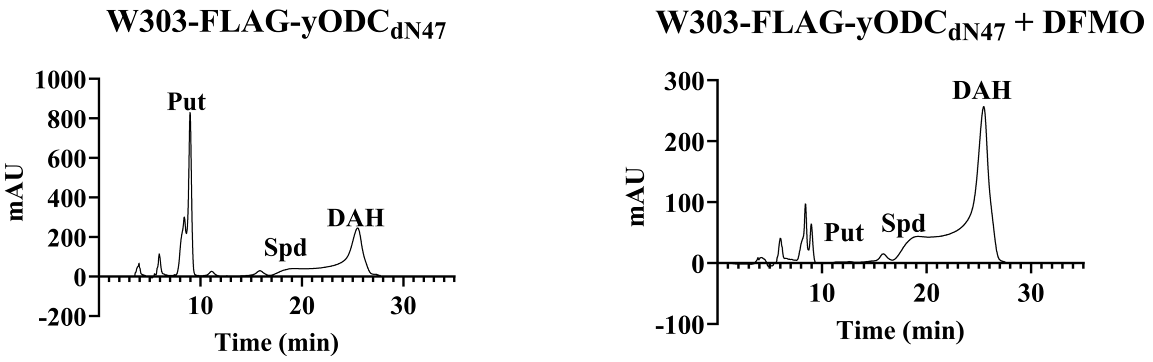


**Supplementary Figure 7**. Representative HPLC spectra for polyamine samples of Figure 2h.

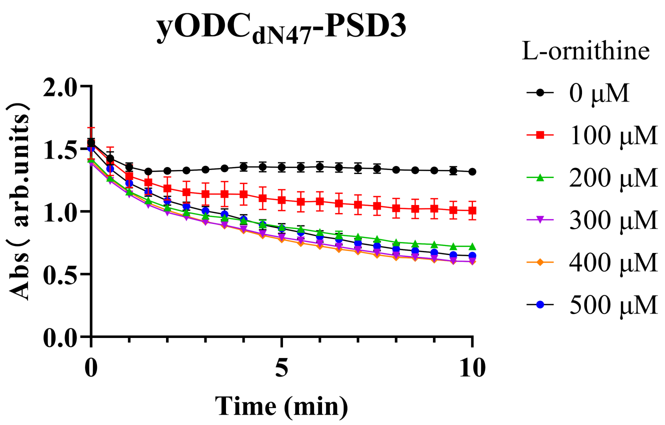


**Supplementary Figure 8**. Enzymatic activity data of yODC$_{dN47}$-PSD3 from the $CO_2$ detection kit.

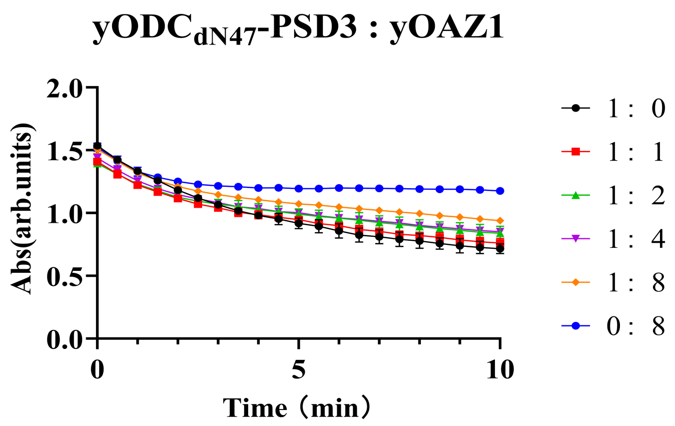


**Supplementary Figure 9**. Inhibition of yODC$_{dN47}$-PSD3 activity by yOAZ1. The molar ratios of yODC$_{dN47}$-PSD3 and yOAZ1 are shown in the legends.

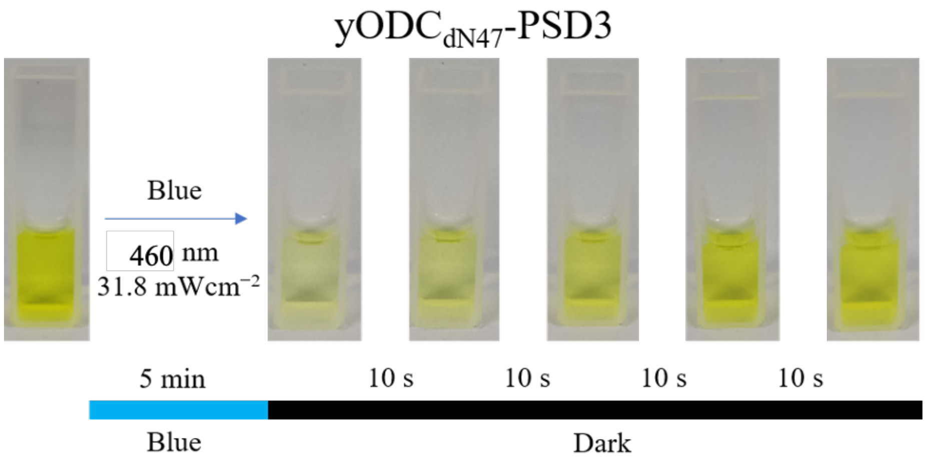


**Supplementary Figure 10**. The color change of the yODC$_{dN47}$-PSD3 protein solution upon light excitation and dark treatment.

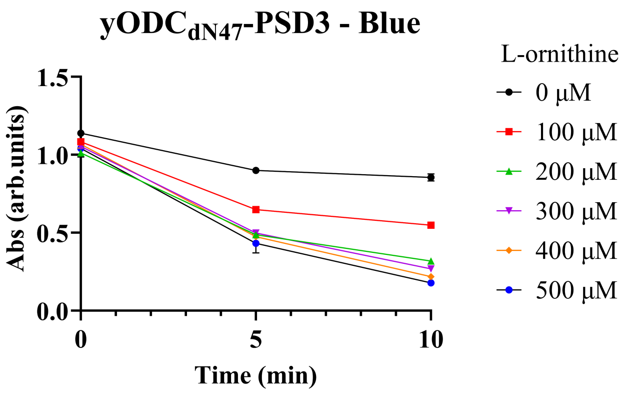


**Supplementary Figure 11**. Enzymatic activity data of yODC$_{dN47}$-PSD3 after light excitation from the $CO_2$ detection kit.

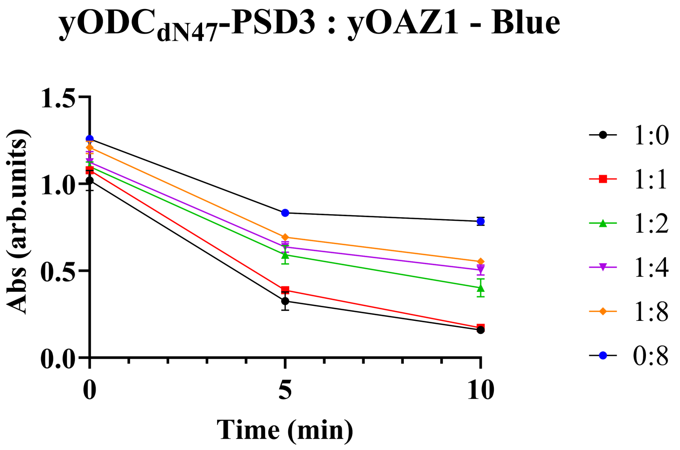


**Supplementary Figure 12**. Inhibition of yODC$_{dN47}$-PSD3 activity by yOAZ1 after light excitation.

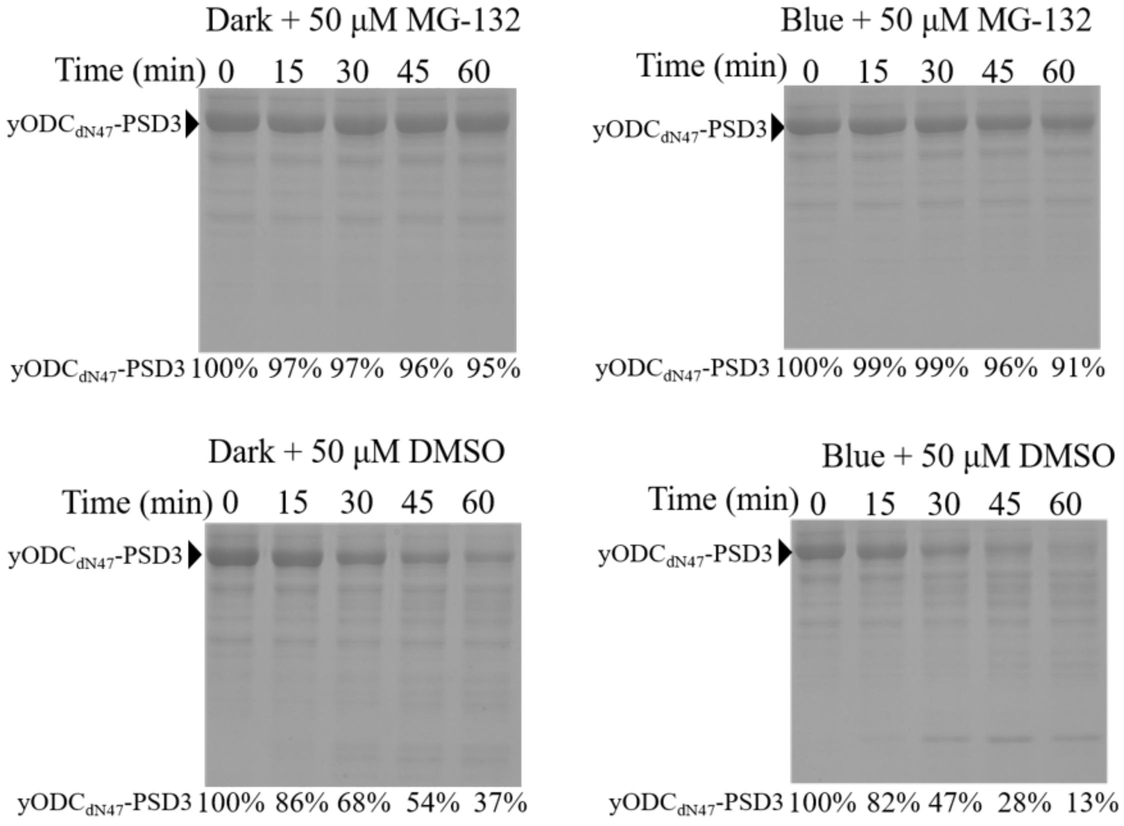


**Supplementary Figure 13**. The full SDS-PAGE gel of Figure 3g.

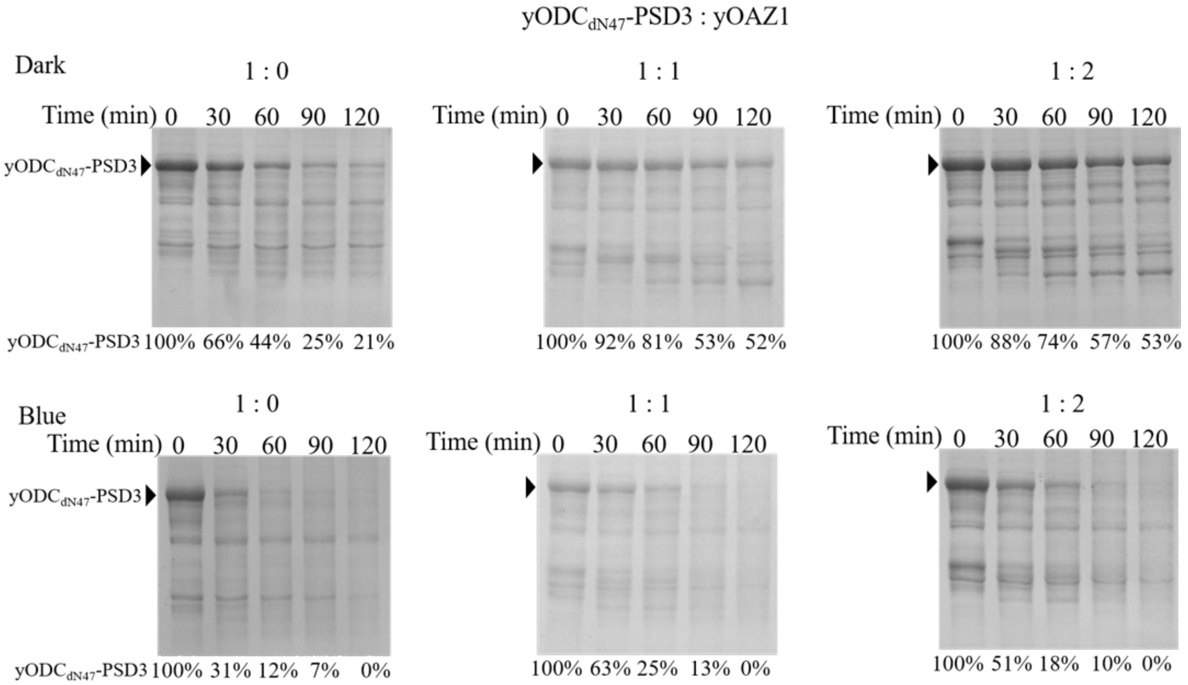


**Supplementary Figure 14**. The full SDS-PAGE gel of Figure 3h.

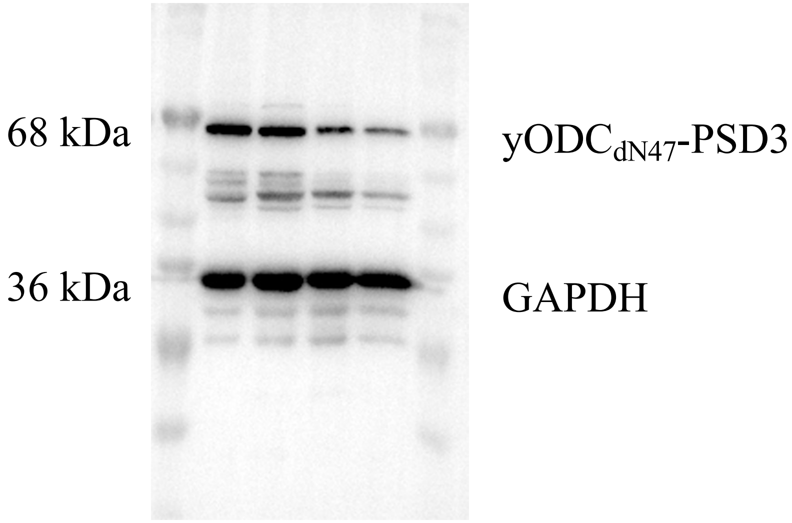


**Supplementary Figure 15**. The full western blotting gel of Figure 4c.

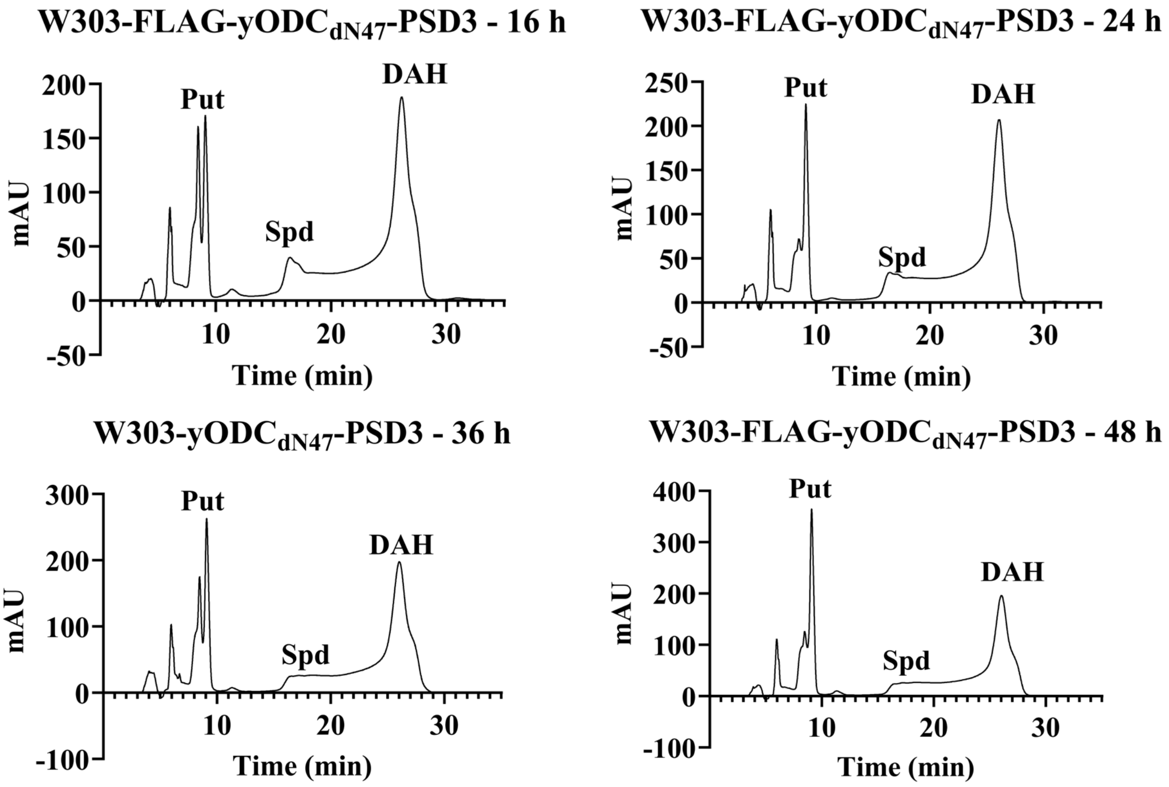


**Supplementary Figure 16**. Representative HPLC spectra of polyamines in W303-FLAG-yODC$_{dN47}$-PSD3 shown in Figure 4d.

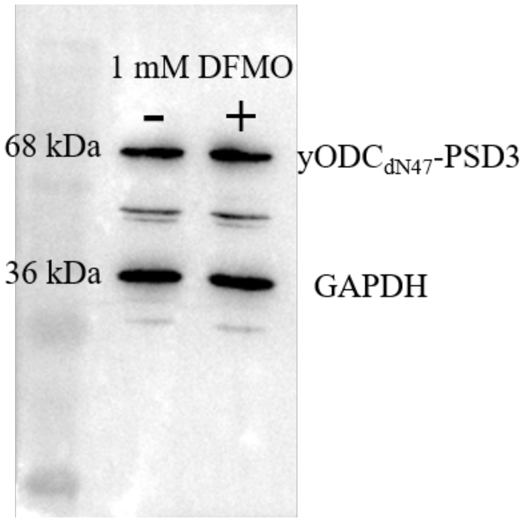


**Supplementary Figure 17**. The full western blotting gel of Figure 4f.

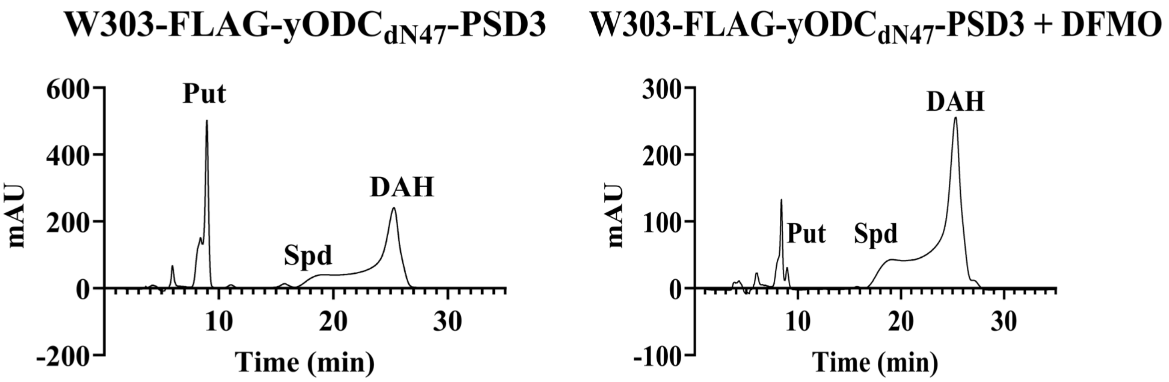


**Supplementary Figure 18**. Representative HPLC spectra of polyamines in W303-FLAG-yODC$_{dN47}$-PSD3 shown in Figure 4g.

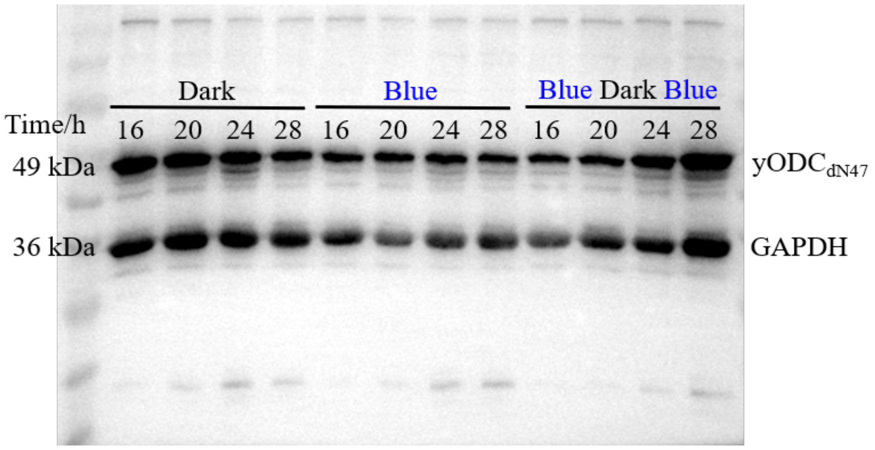


**Supplementary Figure 19**. The full western blotting gel of Figure 5a.

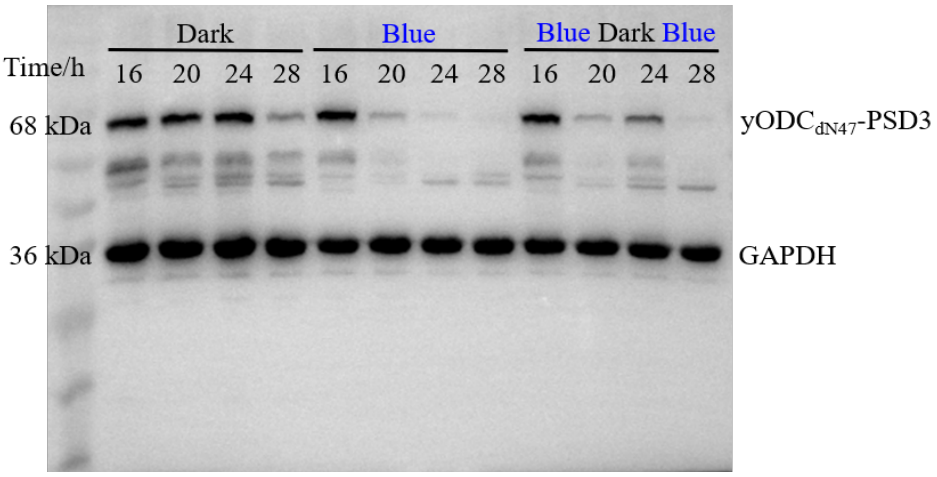


**Supplementary Figure 20**. The full western blotting gel of Figure 5b.

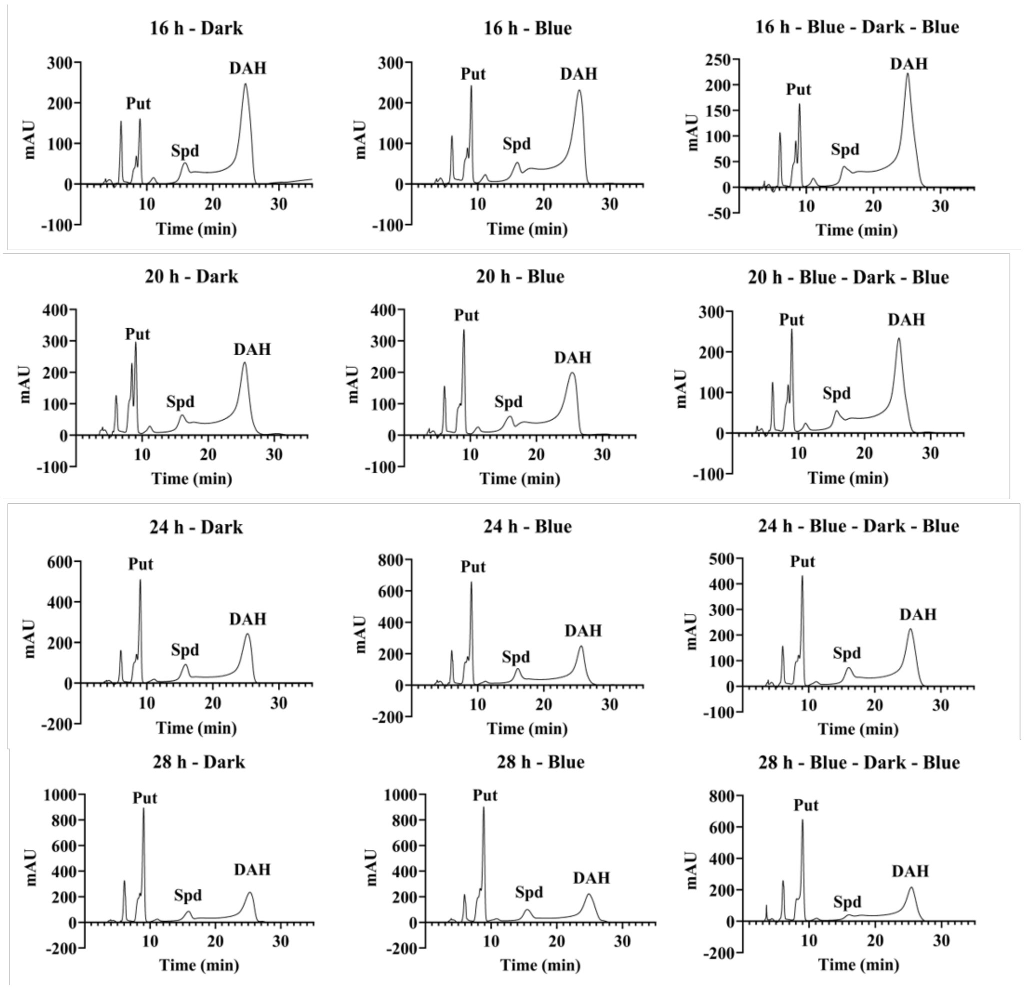


**Supplementary Figure 21**. Representative HPLC spectra of polyamines in W303-FLAG-yODC$_{dN47}$ shown in Figure 5c.

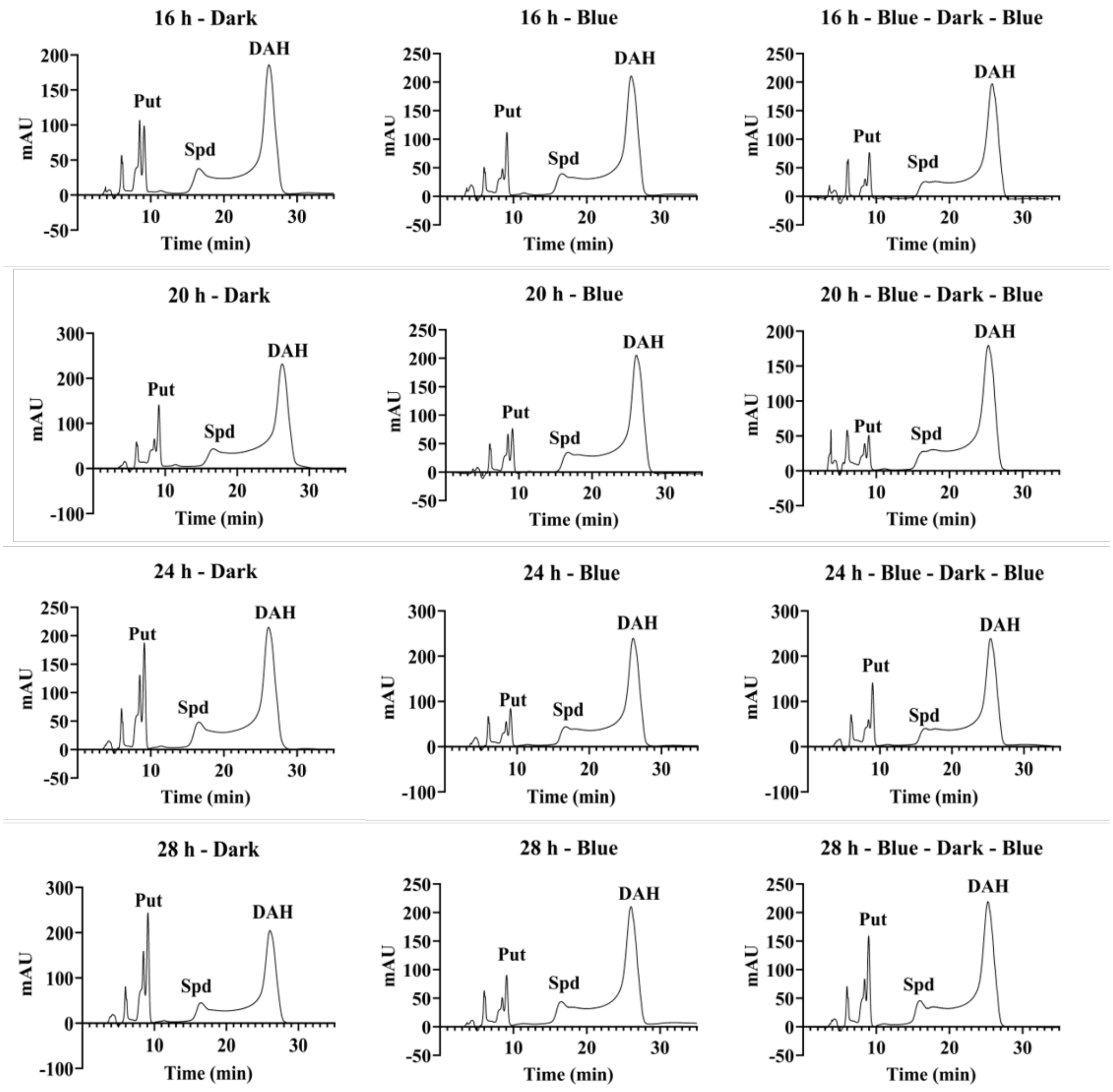


**Supplementary Figure 22**. Representative HPLC spectra of polyamines in W303-FLAG-yODC$_{dN47}$-PSD3 shown in Figure 5d.

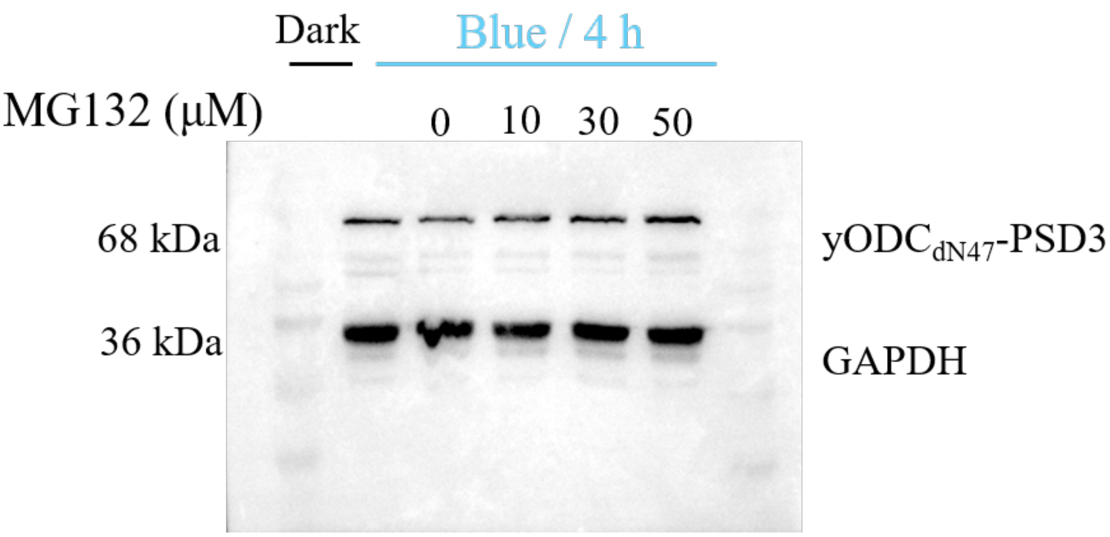


**Supplementary Figure 23**. The full western blotting gel of Figure 5e.

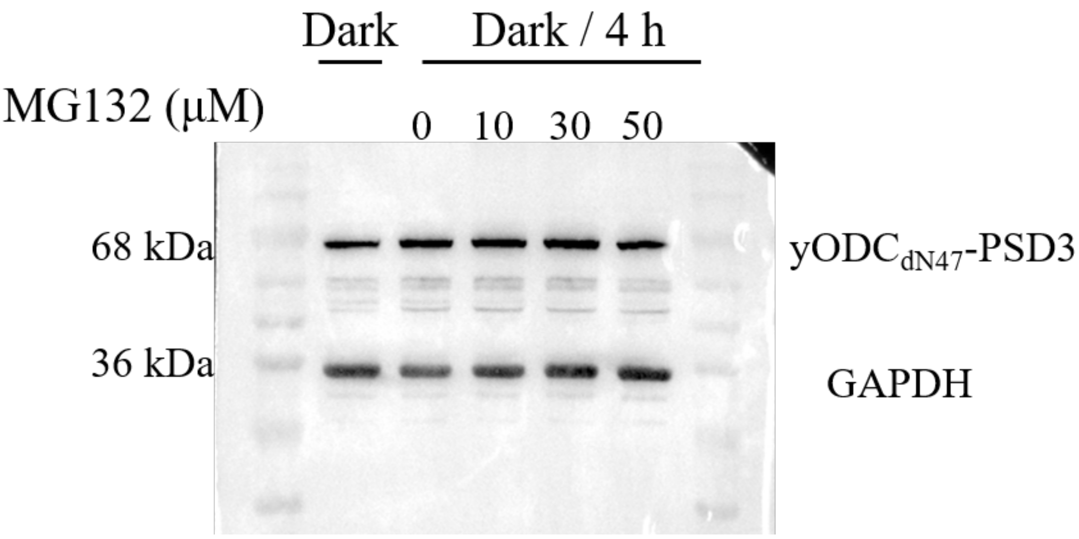


**Supplementary Figure 24**. The full western blotting gel of Figure 5f.

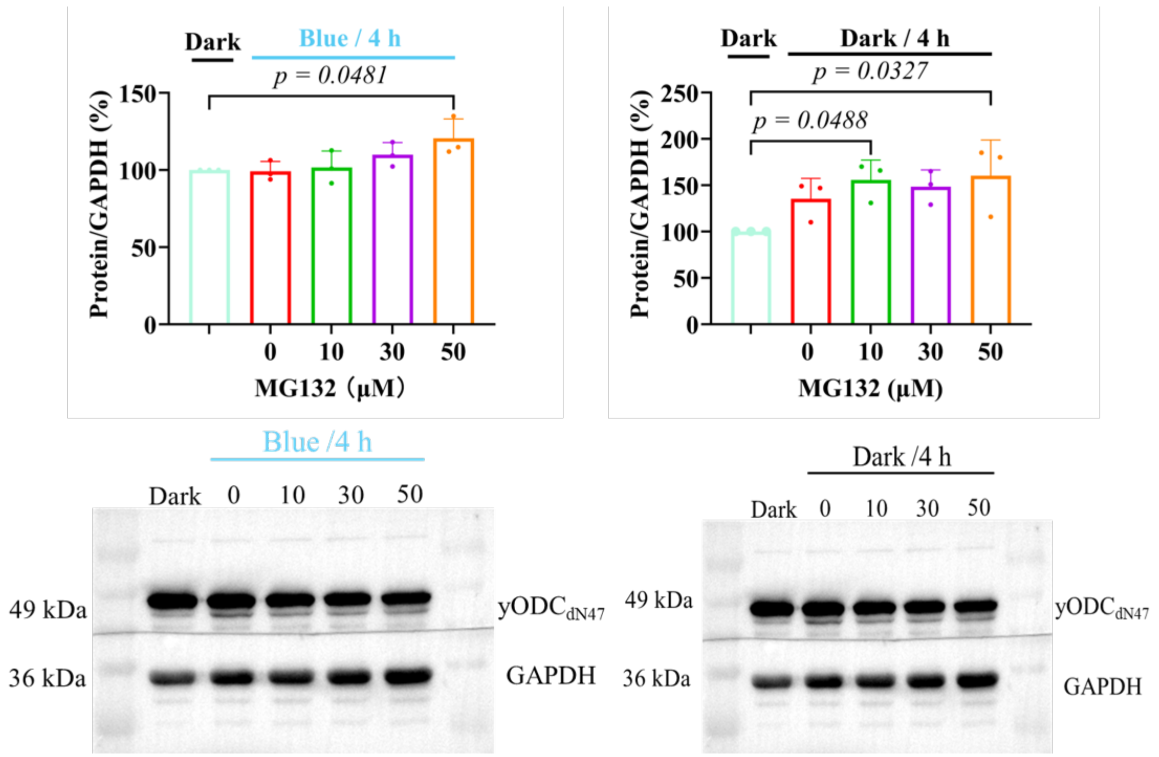


**Supplementary Figure 25**. yODC$_{dN47}$ in W303-FLAG-yODC$_{dN47}$ remained accumulating in both constant darkness and constant light with the existence of MG132. Representative western blotting gels are shown under the statistical data from three biological replicates (mean±s.d., n=3).

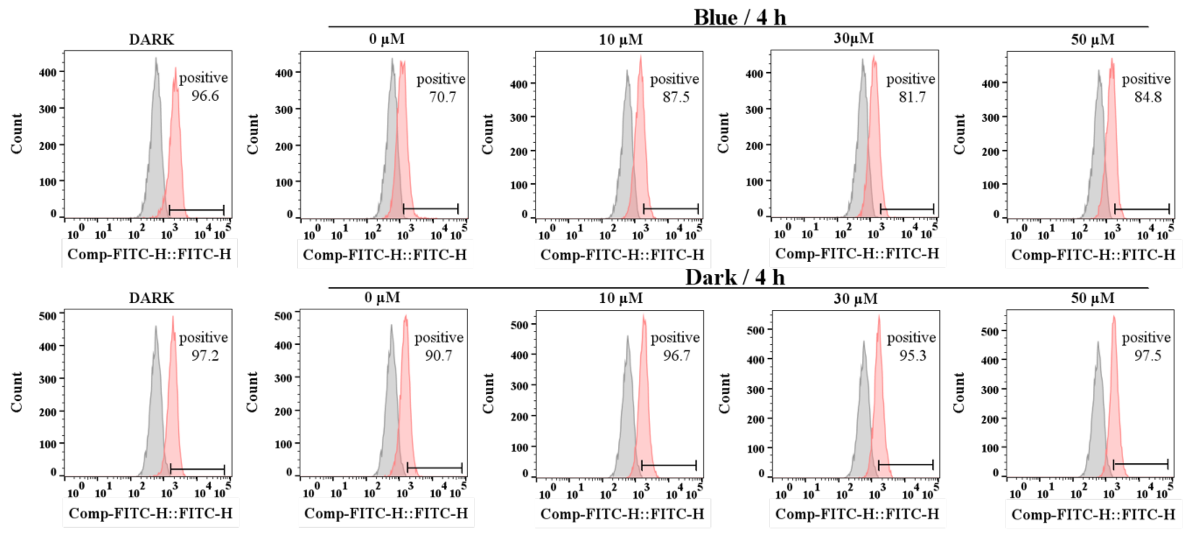


**Supplementary Figure 26**. Representative ROS data of W303-FLAG-yODC$_{dN47}$-PSD3 shown in Figure 5g and Figure 5h.

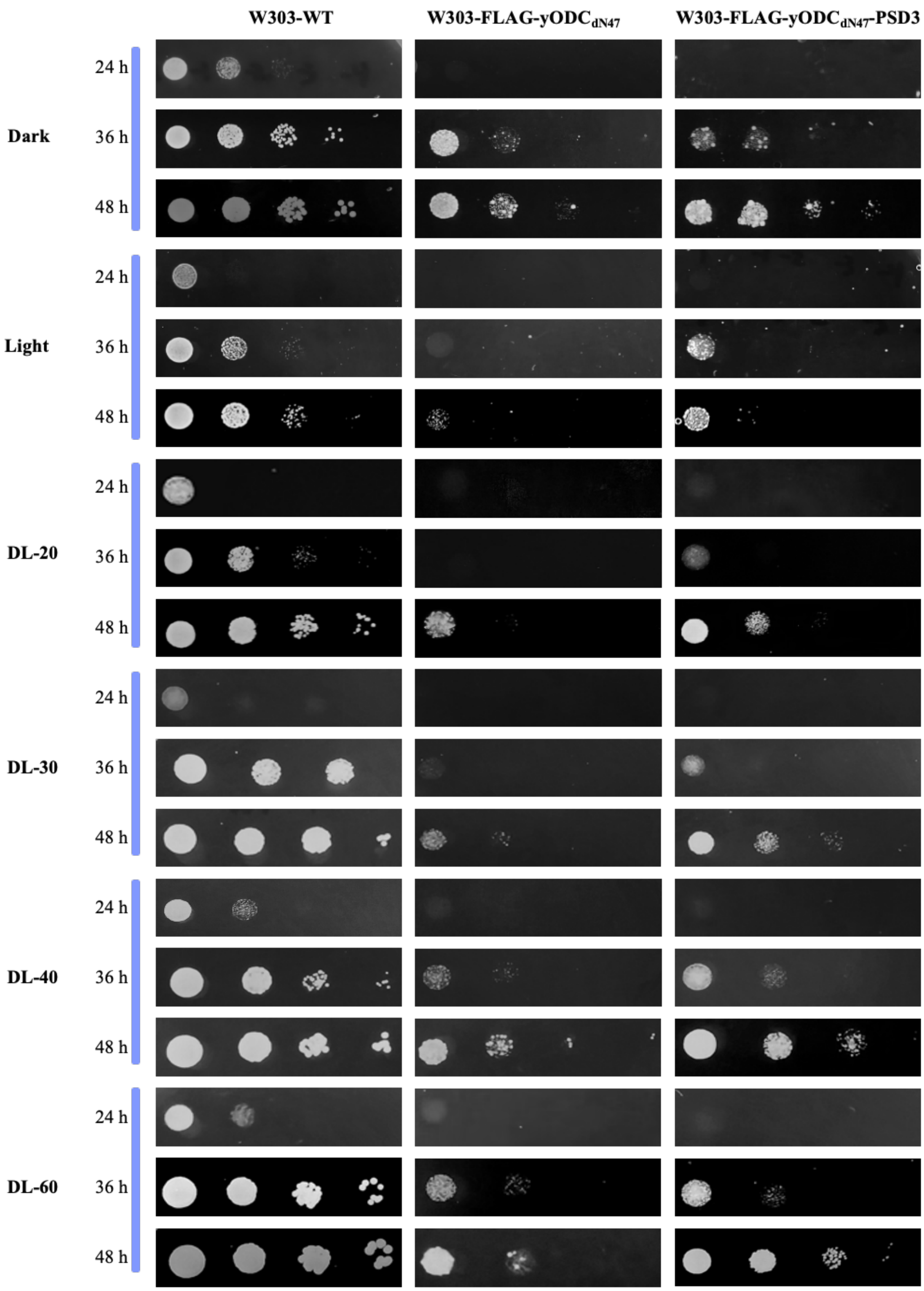


**Supplementary Figure 27**. Growth of different yeast cells under various darkness/light conditions.

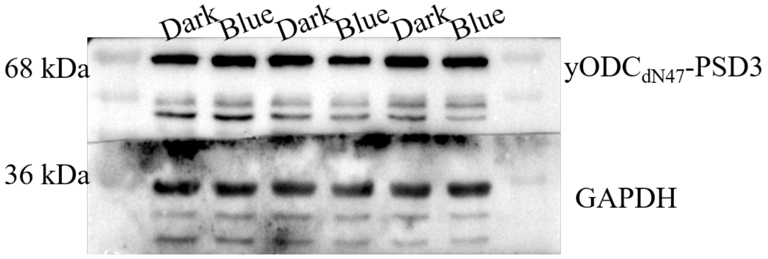


**Supplementary Figure 28**. The full western blotting gel of Figure 6b.

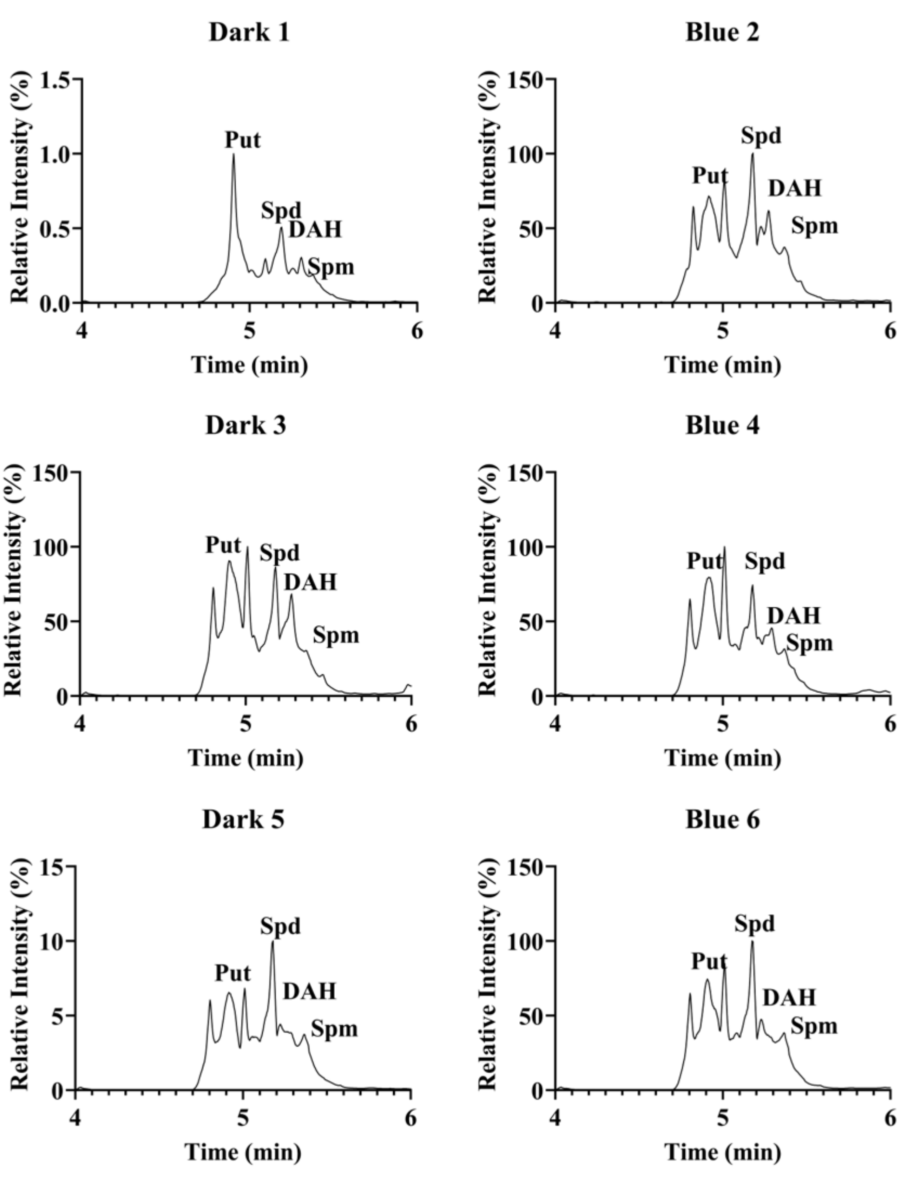


**Supplementary Figure 29**. Representative HPLC spectra of polyamines in W303-FLAG-yODC$_{dN47}$-PSD3 shown in Figure 6c.

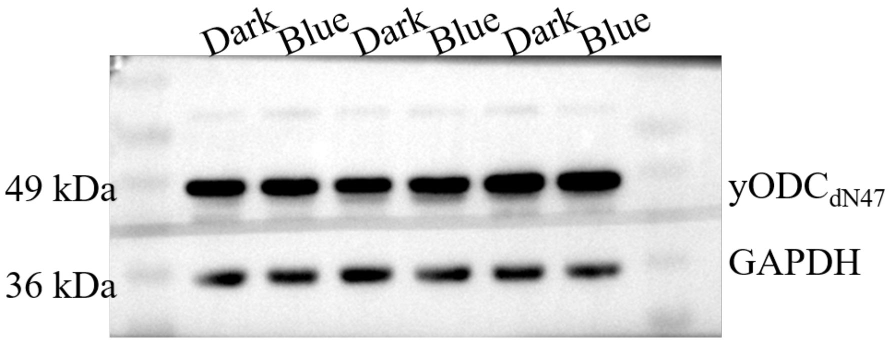


**Supplementary Figure 30**. A representative western blotting gel of Figure 6d.

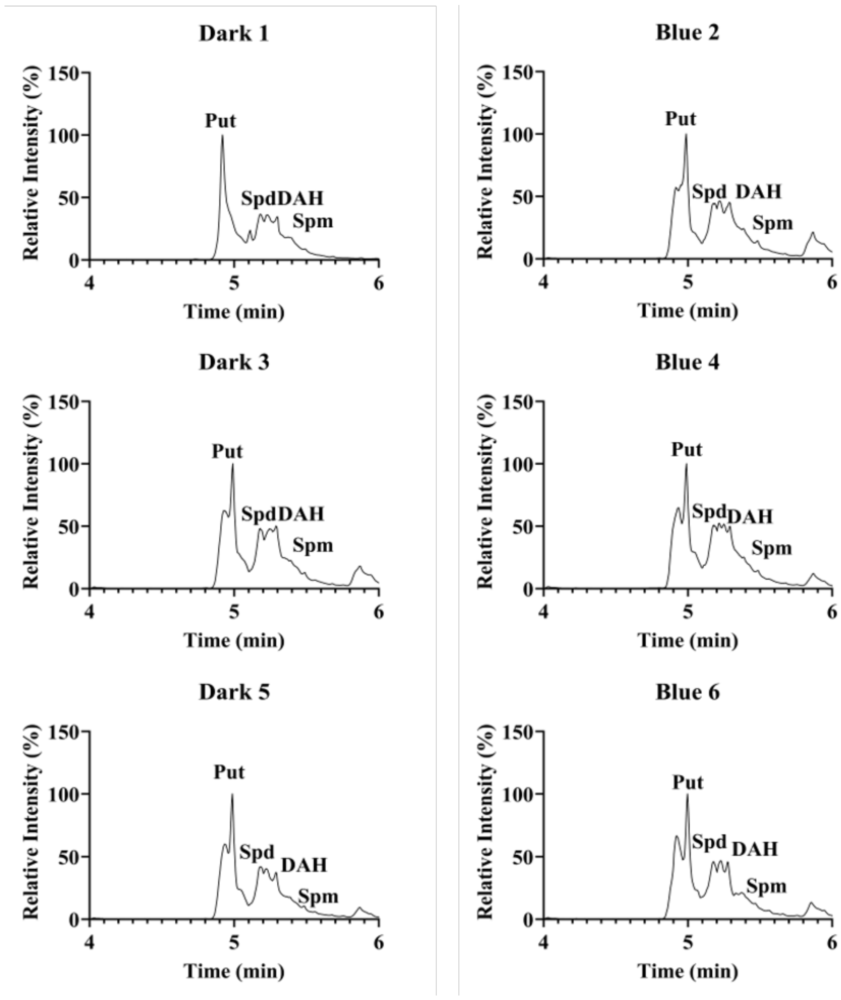


**Supplementary Figure 31**. Representative HPLC spectra of polyamines in W303-FLAG-yODC$_{dN47}$ shown in Figure 6e.

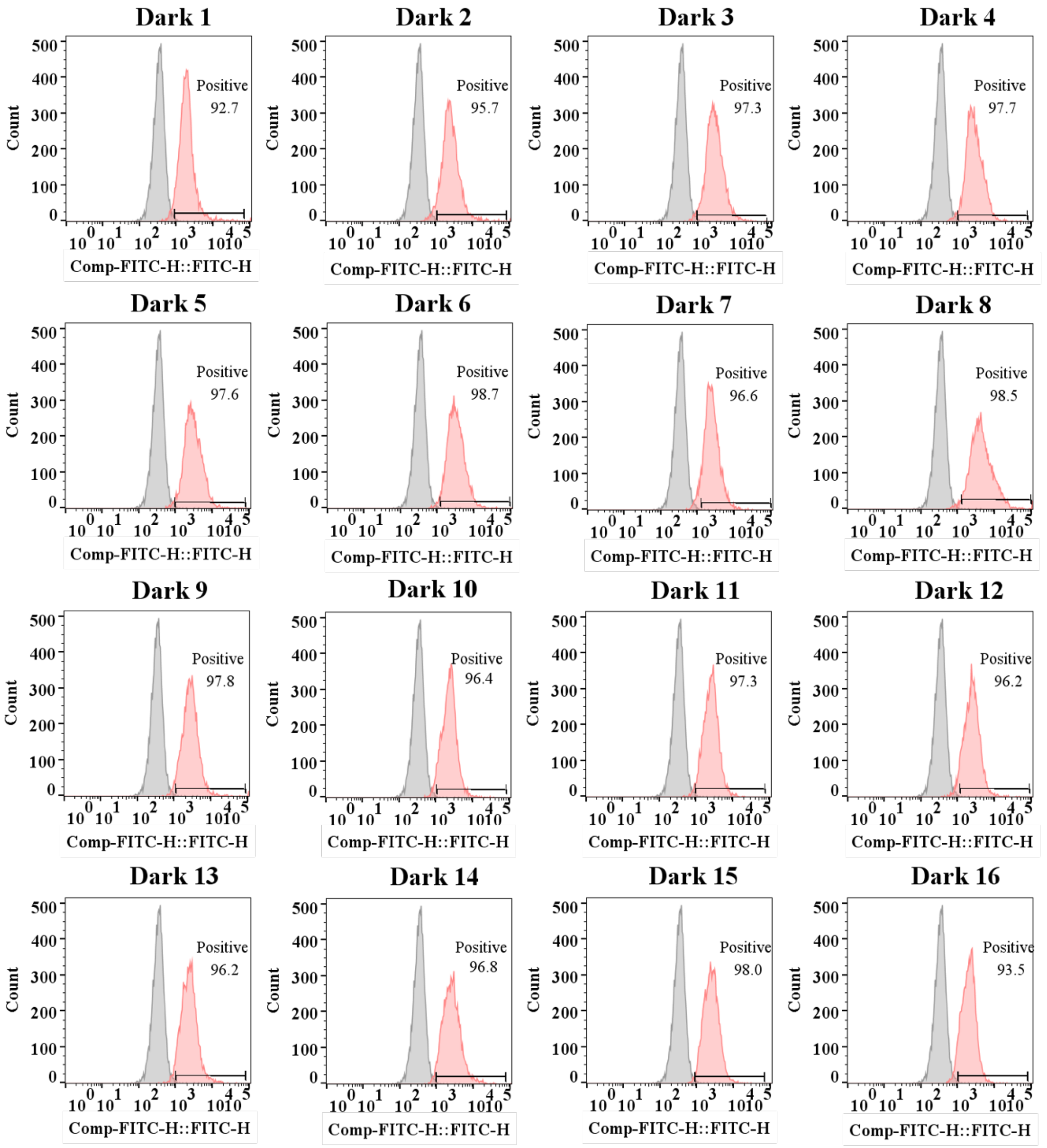


**Supplementary Figure 32**. Representative ROS data of W303-FLAG-yODC$_{dN47}$-PSD3 in constant darkness. The data were collected every 40 min.

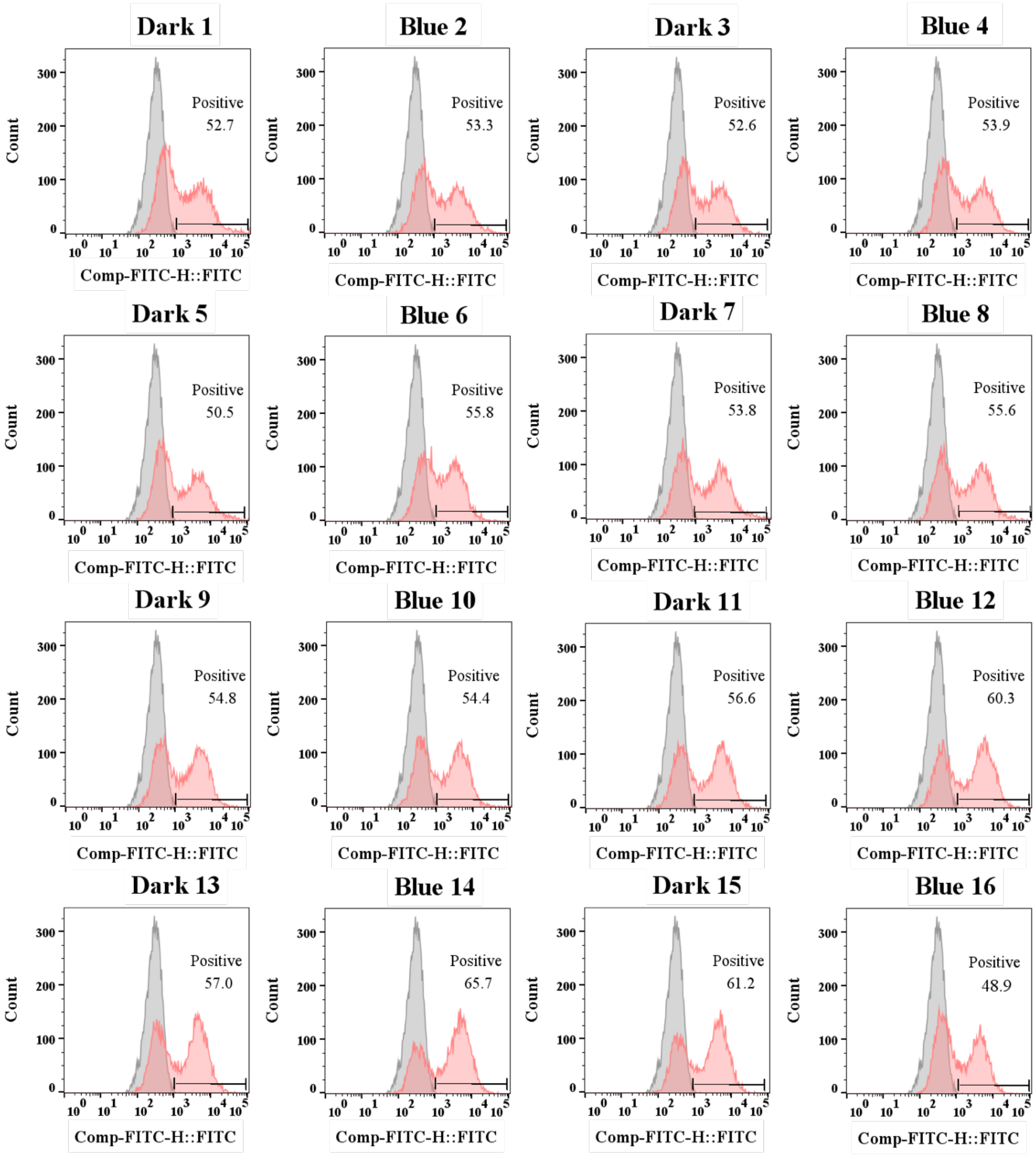


**Supplementary Figure 33**. Representative ROS data of W303-FLAG-yODC$_{dN47}$-PSD3 under DL-40. The data were collected every 40 min.

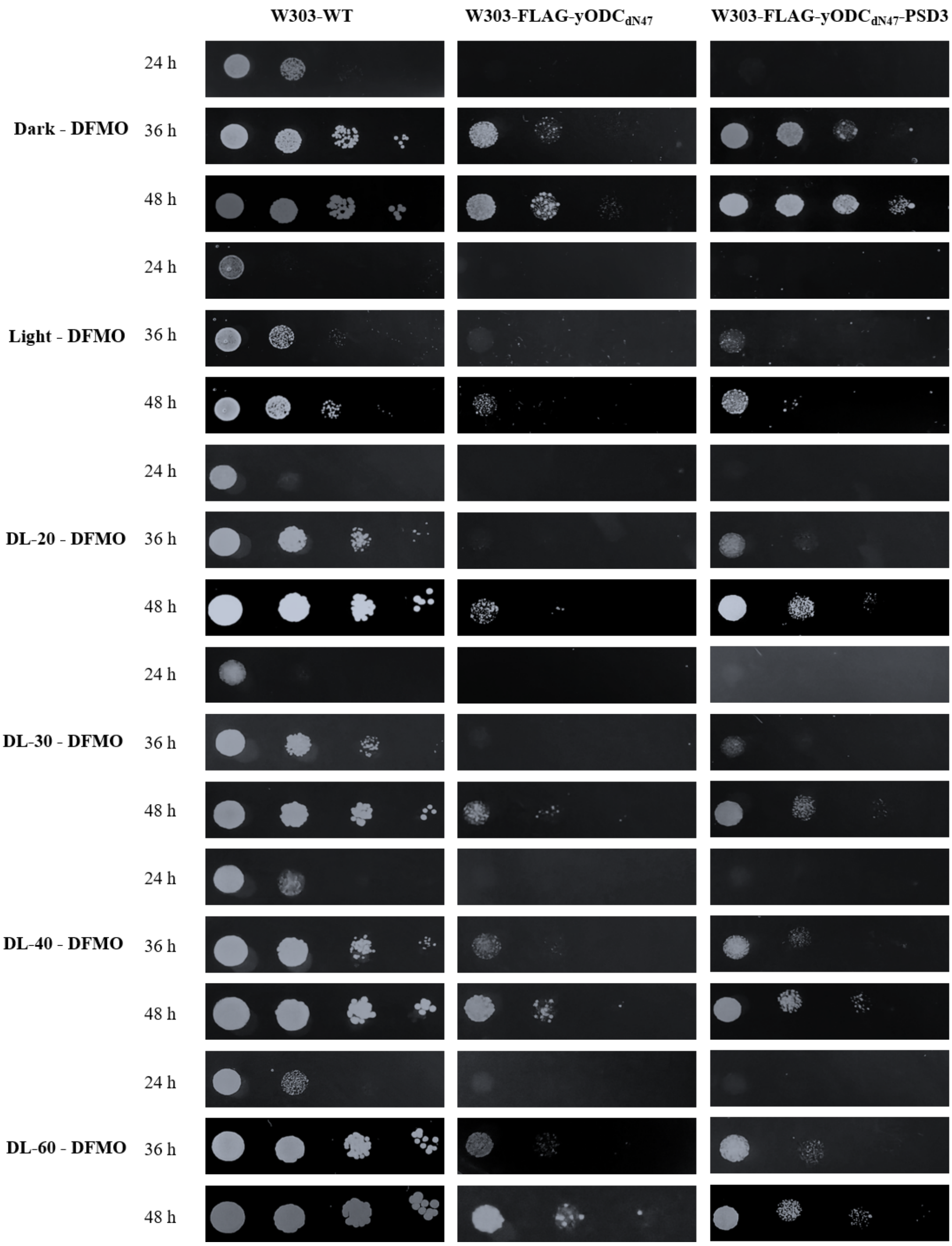


**Supplementary Figure 34**. Growth of different yeast cells under various darkness/light conditions with DFMO.

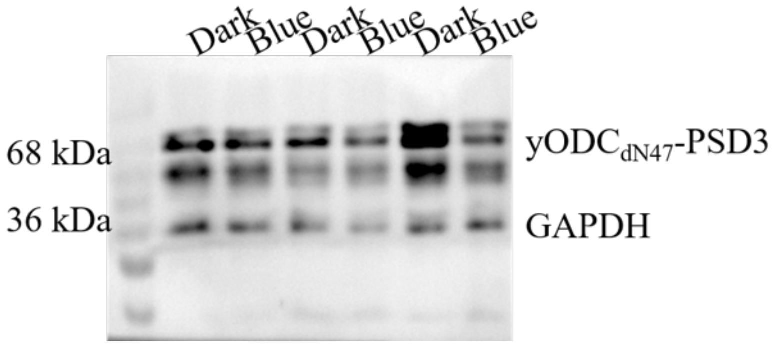


**Supplementary Figure 35**. The full western blotting gel of Figure 7c.

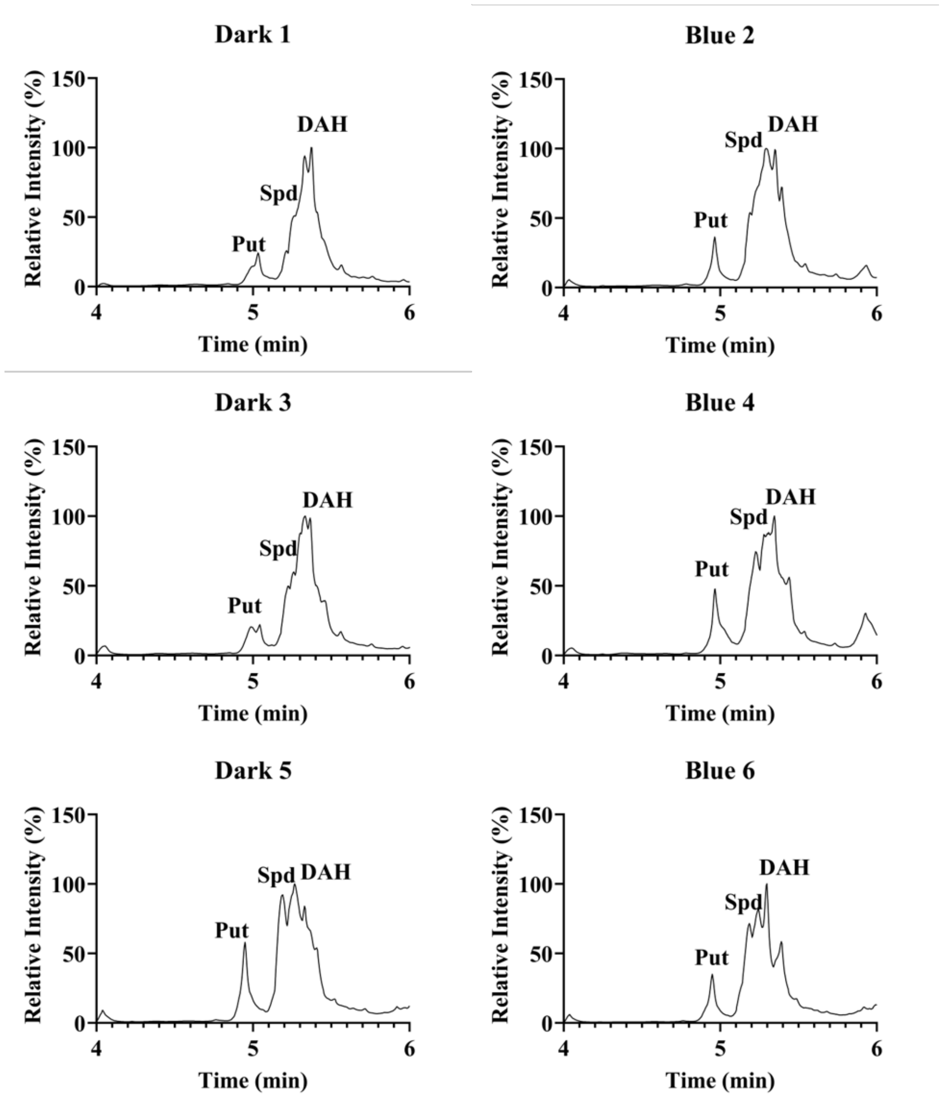


**Supplementary Figure 36**. Representative HPLC spectra of polyamines in W303-FLAG-yODC$_{dN47}$-PSD3 shown in Figure 7d.

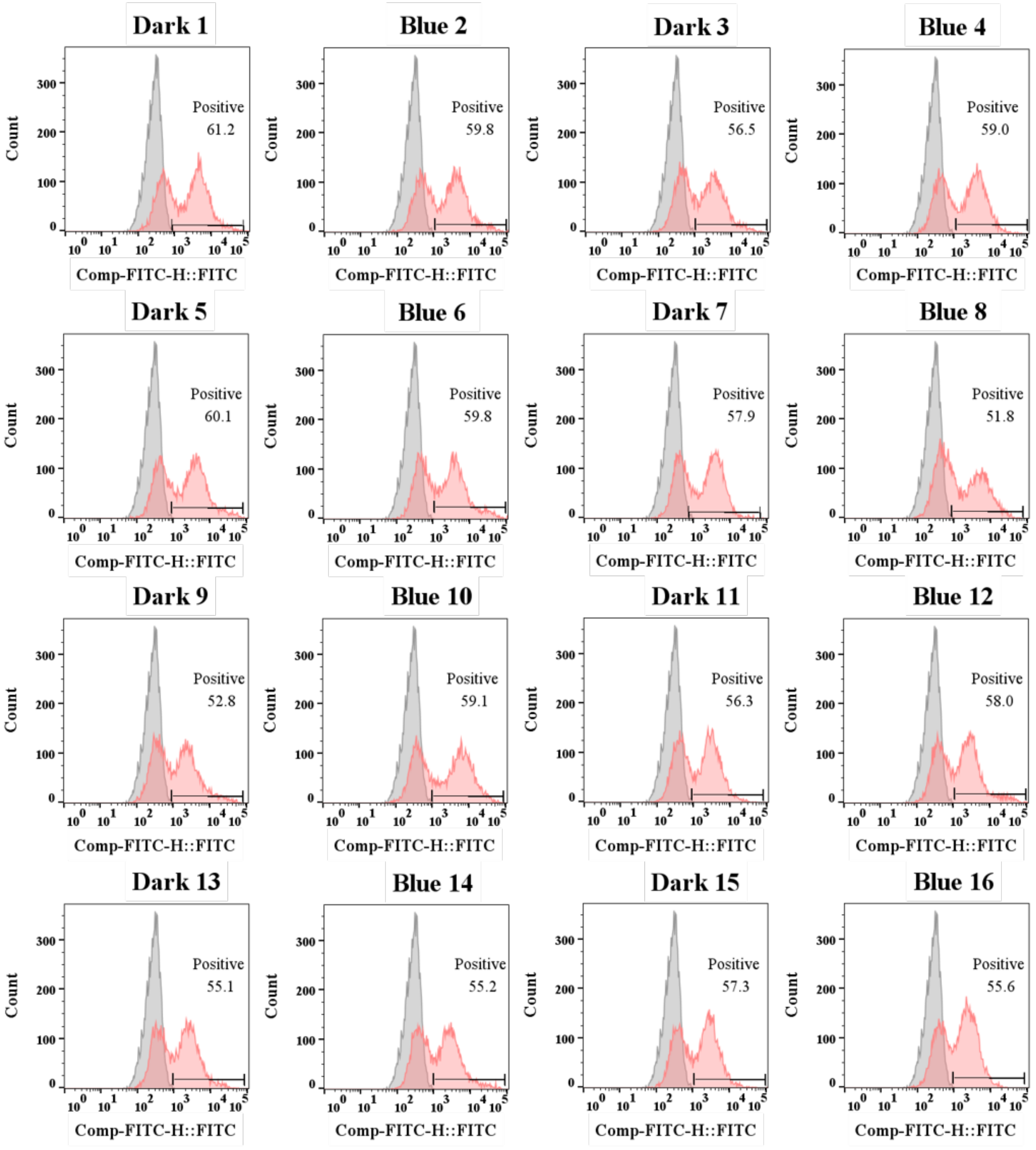


**Supplementary Figure 37**. Representative ROS data of W303-FLAG-yODC$_{dN47}$-PSD3 under DL-40 and DFMO treatment.

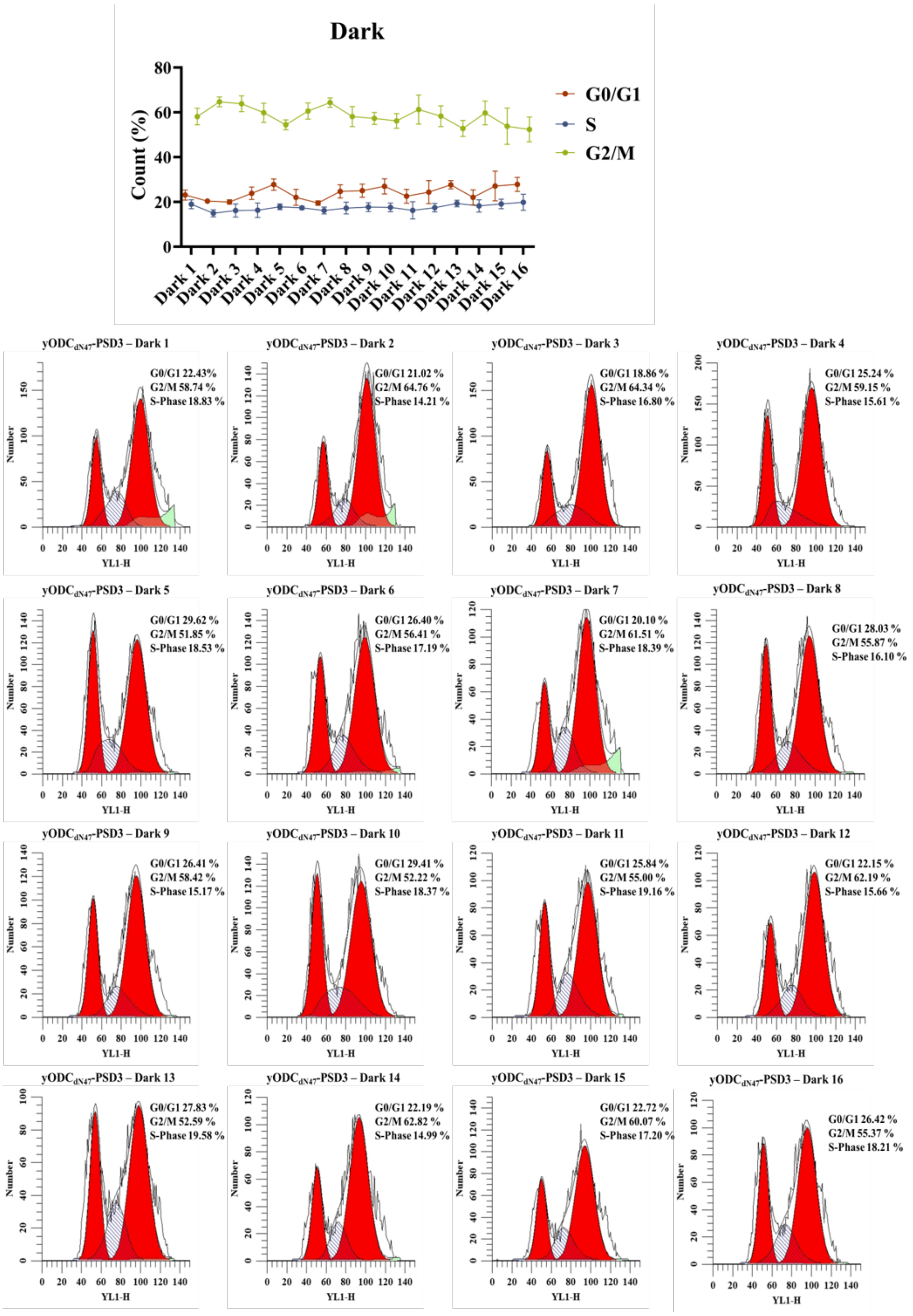


**Supplementary Figure 38**. Representative cell cycle data of W303-FLAG-yODC$_{dN47}$-PSD3 in darkness. Samples were collected every 40 min, and 10,000 cells were counted for each sample. The percentages of the cell numbers of different cell cycle phases from three biological replicates were shown in the top statistical figure (mean±s.d.).

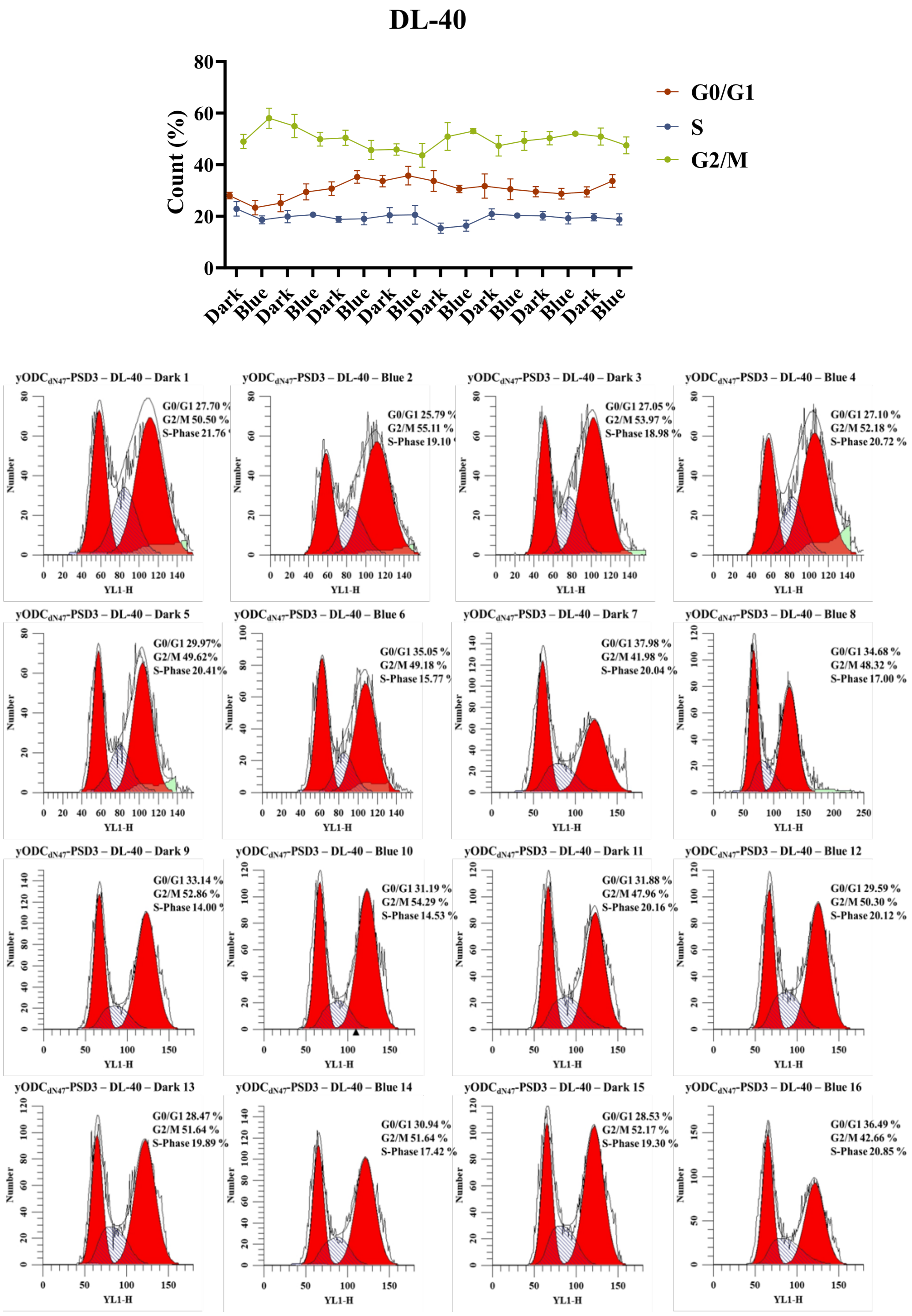


**Supplementary Figure 39**. Representative cell cycle data of W303-FLAG-yODC$_{dN47}$-PSD3 under DL-40. Samples were collected every 40 min, and 10,000 cells were counted for each sample. The percentages of the cell numbers of different cell cycle phases from three biological replicates were shown in the top statistical figure (mean±s.d.).

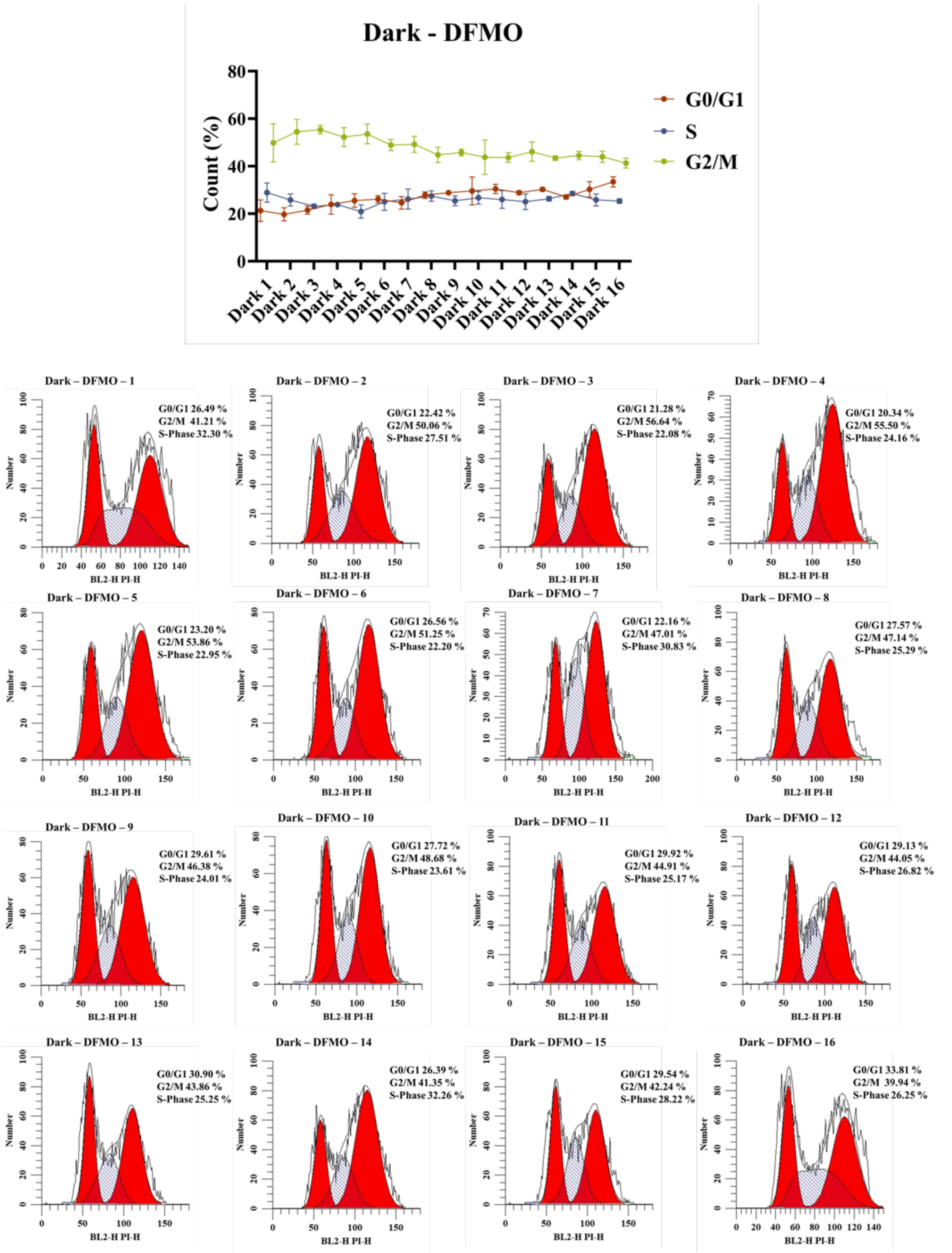


**Supplementary Figure 40**. Representative cell cycle data of W303-FLAG-yODC$_{dN47}$-PSD3 in darkness and DFMO (1mM). Samples were collected every 40 min, and 10,000 cells were counted for each sample. The percentages of the cell numbers of different cell cycle phases from three biological replicates were shown in the top statistical figure (mean±s.d.).

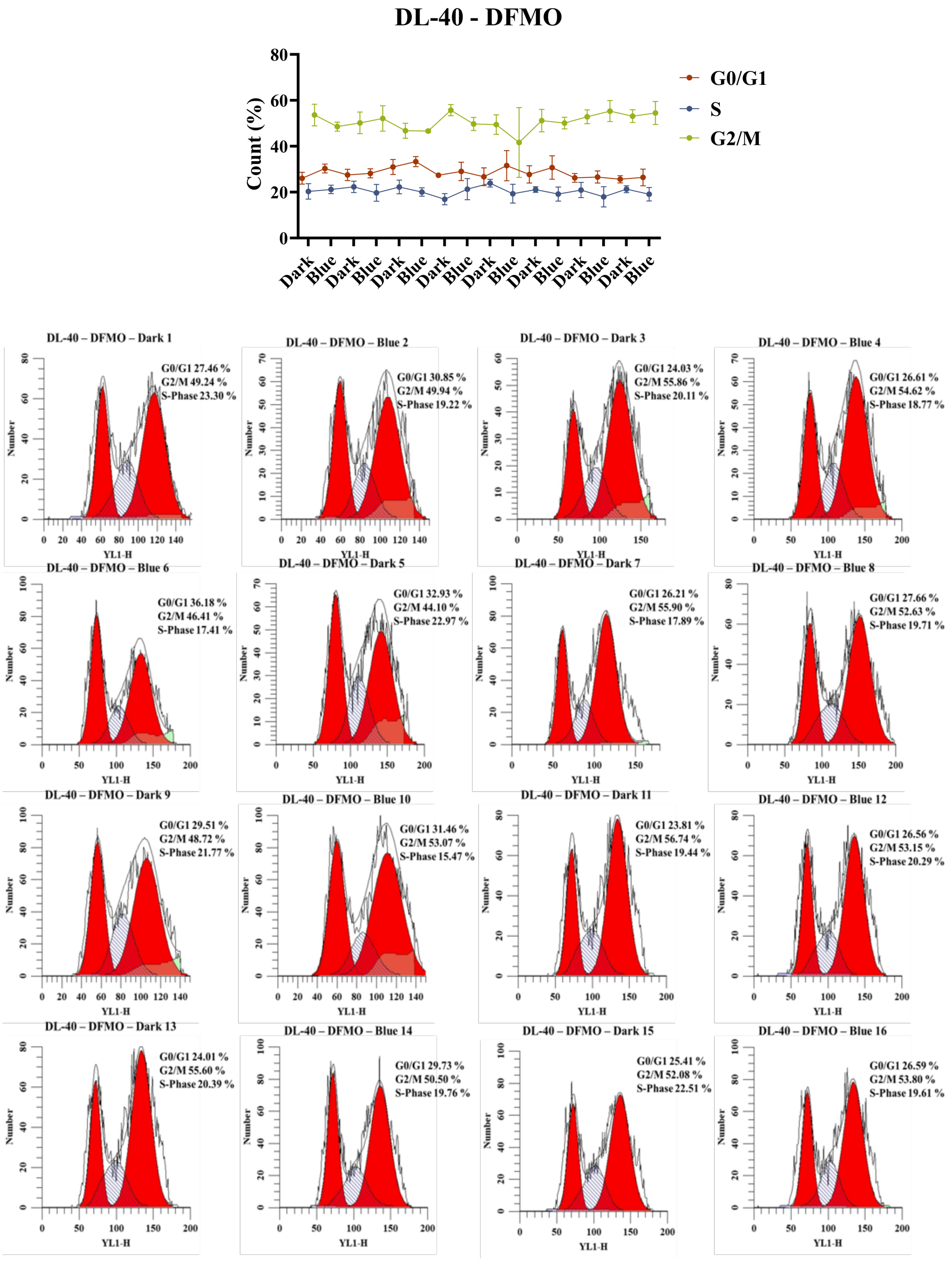


**Supplementary Figure 41**. Representative cell cycle data of W303-FLAG-yODC$_{dN47}$-PSD3 under DL-40 and DFMO (1mM). Samples were collected every 40 min, and 10,000 cells were counted for each sample. The percentages of the cell numbers of different cell cycle phases from three biological replicates were shown in the top statistical figure (mean±s.d.).
